# Genome-edited retinal organoids restore host bipolar connectivity in the primate macula

**DOI:** 10.1101/2025.08.21.671525

**Authors:** Atsuta Ozaki, Akihiro Kawai, Ryutaro Akiba, Satoko Okayama, Nobuhiko Ohno, Keisuke Kajita, Tomohiro Masuda, Satoshi Yokota, Shin-ichiro Ito, Du Peiyan, Kenta Onoue, Shigenobu Yonemura, Mineo Kondo, Yasuo Kurimoto, Yingbin Fu, Michiko Mandai

## Abstract

Retinal organoids (ROs) represent a promising regenerative strategy for restoring vision in retinal degenerative diseases, but whether host cone bipolar cells (BCs) in the primate macula can rewire with transplanted photoreceptors remains unresolved. Here, we transplanted genome-edited human retinal organoids lacking ON-BCs (*Islet-1⁻/⁻* ROs) into a non-human primate macular degeneration model. Remarkably, host rod and cone BCs extended dendrites toward grafted photoreceptors, forming functional synapses confirmed by immunohistochemistry, ultrastructural imaging, and focal macular electroretinography. Both ON- and OFF-pathway connectivity was rebuilt, providing the first demonstration of host–graft synaptic integration in the primate macula. These results establish that primate cone circuits retain a surprising capacity for rewiring and highlight genome-edited ROs as a powerful platform for vision restoration. Our findings represent a critical translational step toward stem cell–based therapies capable of repairing central vision in patients with advanced macular degeneration.

## Introduction

Inherited retinal diseases and age-related macular degeneration cause irreversible vision loss through progressive photoreceptor degeneration (1, 2). Loss of photoreceptor cells (PRCs) in the macula directly impairs central vision, profoundly affecting quality of life. Stem cell– derived retinal organoids (ROs) represent a promising regenerative strategy (3–5), but true restoration of vision will depend on whether host BCs can re-establish synaptic connections with transplanted photoreceptors. This requirement is particularly critical for cone BCs in the primate macula, where cone circuits dominate central vision.

Rodent transplantation studies have shown that developing ROs can integrate with host retina, form synapses with rod BCs, and restore light responses (3, 6–8). Recent clinical study has further demonstrated the safety and long-term survival of organoid grafts in patients (9). However, direct evidence that primate cone BCs can regenerate dendrites and form functional synapses with donor PRCs is lacking. Rodent models are limited by the absence of a macula, and prior non-human primate (NHP) transplants were placed outside the fovea, preventing assessment of cone pathway rewiring.

Previous works suggest that BC plasticity varies by subtype and declines with age (10–12). Rod BC dendritic outgrowth has been observed in degenerative conditions (13–15), but cone BC plasticity remains uncertain. In primates, a recent study demonstrated that midget BCs in the fovea of zones with complete PRC degeneration failed to extend dendrites toward the neighboring remaining PRCs, raising doubts about their regenerative capacity (16). Conversely, transplantation of developing ROs into rodent retinas has been shown to stimulate BC dendritic regrowth and recruitment of horizontal cells, suggesting that a developmental graft environment may enhance host plasticity (17, 18). Whether such plasticity extends to cone BCs in the primate macula, a key requirement for restoring cone-mediated central vision, has remained unresolved.

Here, we address this gap by transplanting genome-edited retinal organoid sheets derived from human embryonic stem cells lacking ON-BCs (*Islet-1⁻/⁻*) into a NHP model of macular degeneration. Removing graft-derived ON-BCs eliminates competition for synaptic partners and enables direct assessment of host-driven rewiring (6, 7). We demonstrate that primate rod and cone BCs extend dendrites into grafts, form canonical synapses with transplanted photoreceptors, and support recovery of b-wave responses in focal macular electroretinography. These findings provide the first demonstration of functional host–graft synaptic integration in the primate macula and highlight the potential of genome-edited ROs as a strategy to restore cone-mediated central vision in macular degeneration.

## Results

### Generation of an NHP Macular Degeneration Model via Parafoveal Laser Photocoagulation

To examine synaptic connections between different cone BC subtypes and graft PRCs, we induced retinal degeneration via laser photocoagulation in the parafovea and extrafoveal regions, including areas with the highest cone-BC density (19) (Supplemental Figure 1A). Based on our previous report (20), We first optimized the laser power by testing coagulations outside the macula at power levels ranging from 40mW to 150mW. Fundus images revealed white lesions immediately after laser application, which became less apparent over time at 40mW-120mW, while 150mW laser caused an immediate retinal hemorrhage (Supplemental Figure 1B). Autofluorescence imaging showed initial hyperfluorescence at all laser powers, followed by hypofluorescence (Supplemental Figure 1B). Optical coherence tomography (OCT) imaging demonstrated consistent diffuse opacification in the outer nuclear layer (ONL) immediately after laser coagulation at 60–80mW, leading to selective ONL loss at one week and one month post-treatment (Supplemental Figure 1C). Higher power levels (100-150mW) also damaged the inner nuclear layer (INL) at one month after treatment (Supplemental Figure 1C).

Based on these findings, we performed laser photocoagulation in the parafovea and perifovea regions using 40mW and 70mW (Supplemental Figure 2A). At one month, autofluorescence imaging showed mottling in the coagulated area, suggesting localized retinal pigment epithelium (RPE) impairment (Supplemental Figure 2A). OCT confirmed that 70-mW laser treatment caused a complete ONL loss at one month, while ONL remained at 40mW (Supplemental Figure 2B). Focal macular electroretinography (FMERG) recording from the 70mW lesions were nearly non-recordable, where those from the 40mW showed moderate b-wave (Supplemental Figure 2C). Laser photocoagulation selectively ablated ONL while preserving INL at laser power of 70mW by histology (Supplemental Figure 2D-I). Rhodopsin [outer segment (OS) of rod PRCs], L/M Opsin (OS of cone PRCs), S Opsin, and Recoverin (PRCs)-positive cells were almost lost with 70mW laser coagulation (Supplemental Figure 2E and F). SCGN [diffuse bipolar (DB)1, 6], Calretinin (amacrine cells), and Calbindin [horizontal cells (HCs), DB3a, 6]-positive cells were similarly observed in the eyes with laser at 40mW, 70mW, and no treatment (Supplemental Figure 2F and G). The number of RPE65-positive cells decreased at 70mW lesions, indicating a potential impact on RPE (Supplemental Figure 2H). Additionally, GFAP expression was increased in the photoreceptor ablated retinas, indicating the activation of Müller cells, mimicking retinal degeneration (Supplemental Figure 2I). Based on these results, laser power of 70mW±10mW was used for further experiments.

### The dendrites of multiple BC subtypes consistently retracted after photocoagulation

Previous reports suggested that BC dendrite length remains unchanged after cone PRC damage caused by laser treatment or diphtheria toxin (21, 22). However, since our model involved near-complete photoreceptor ablation, we assessed BC dendrite length from the INL border to the dendritic tip of different BC subtypes. BCs and HCs were distinguished by the presence of axons extending into the inner plexiform layer (IPL).

Dendrites of EAAT2+ (pan OFF-cone BCs), PKCα+ (DB4, DB5, rod BCs), SCGN+, and Calbindin+ cells showed significant retraction following ONL ablation (Figure 1A-H’), with a marked reduction in dendrite length (Figure 1I-L). We then examined whether retracted dendrites retained postsynaptic markers. mGluR6 immunoreactivity was almost completely lost, consistent with previous reports in laser-treated and genetically degenerated retinas (1,23) (Figure 1M-N’). Conversely, GluK1, an OFF-cone BC postsynaptic marker, remained detectable, though its expression was reduced (Figure 1O-P’).

**Figure 1.**
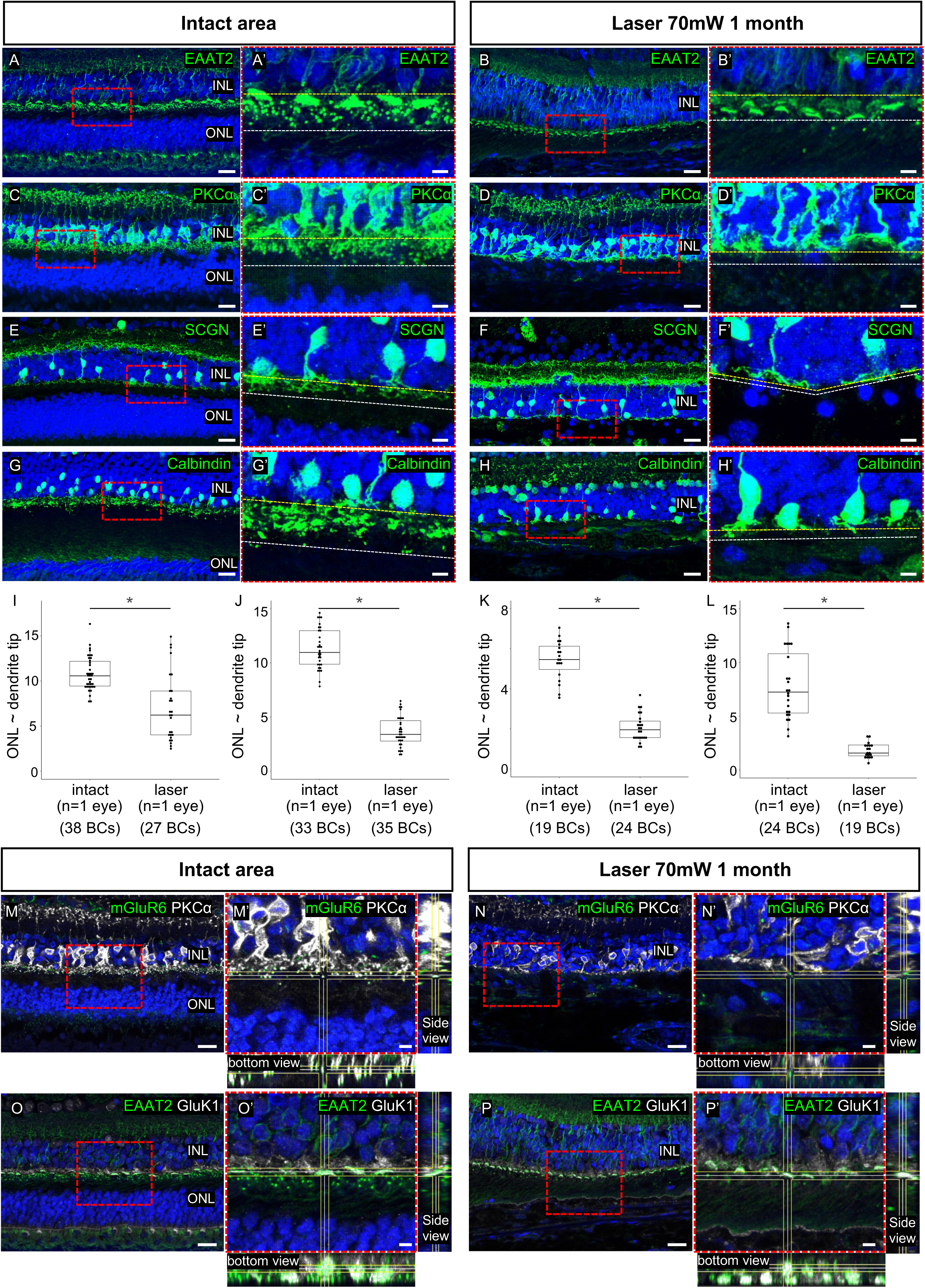
Changes in BC dendrites following laser treatment. (A–H’) Immunohistological analysis of dendritic changes in four BC subtypes one month after laser treatment compared to intact areas. Representative images show EAAT2-positive BCs (A–B’), PKCα-positive BCs (C– D’), SCGN-positive BCs (E–F’), and Calbindin-positive BCs (G–H’). The yellow dashed line marks the bottom of the ONL, the white dashed line indicates dendritic tips, and the red dashed rectangle highlights the enlarged region. (I–L) Quantitative analysis of dendritic length for EAAT2-positive BCs (I), PKCα-positive BCs (J), SCGN-positive BCs (K), and Calbindin-positive BCs (L). Statistical significance was assessed using the Wilcoxon rank sum test (*p < 0.05). (M– N’) Changes in mGluR6-positive postsynaptic markers between intact and laser-treated areas. (O– P’) Changes in GluK1-positive postsynaptic markers between intact and laser-treated areas. Scale bars: 20 μm (A, B, C, D, E, F, G, H, M, N, O, P); 5 μm (A’, B’, C’, D’, E’, F’, G’, H’, M’, N’, O’, P’).

### Long-term survival of Islet-1^-/-^ RO sheets after transplantation under optimized immune suppression

We differentiated *Islet-1⁺/⁺* and *Islet-1⁻/⁻* ROs from the *Crx::Venus* human embryonic stem cell line (*KhES-1*) (Supplemental Figure 3A, B). By differentiation day 60 (dd60), both *Islet-1⁺/⁺*and *Islet-1⁻/⁻* ROs exhibited a characteristic continuous neuroepithelial layer positive for retinal progenitor markers CHX10/PAX6 and apical/basal expression of *Crx::Venus/*BRN3, as confirmed by immunohistochemistry (IHC) (Supplemental Figure 3A’- B’’). We transplanted RO sheets into the eyes of six monkeys (Monkeys 1–2: *Islet-1⁺/⁺* RO sheets; Monkeys 3–6: *Islet-1⁻/⁻* RO sheets), with each eye receiving six sheets (∼1 × 1.5 mm). Graft survival was monitored via *Crx::Venus* expression and OCT (Figure 2A-D’’, Supplemental Figure 3C-J’’).

**Figure 2.**
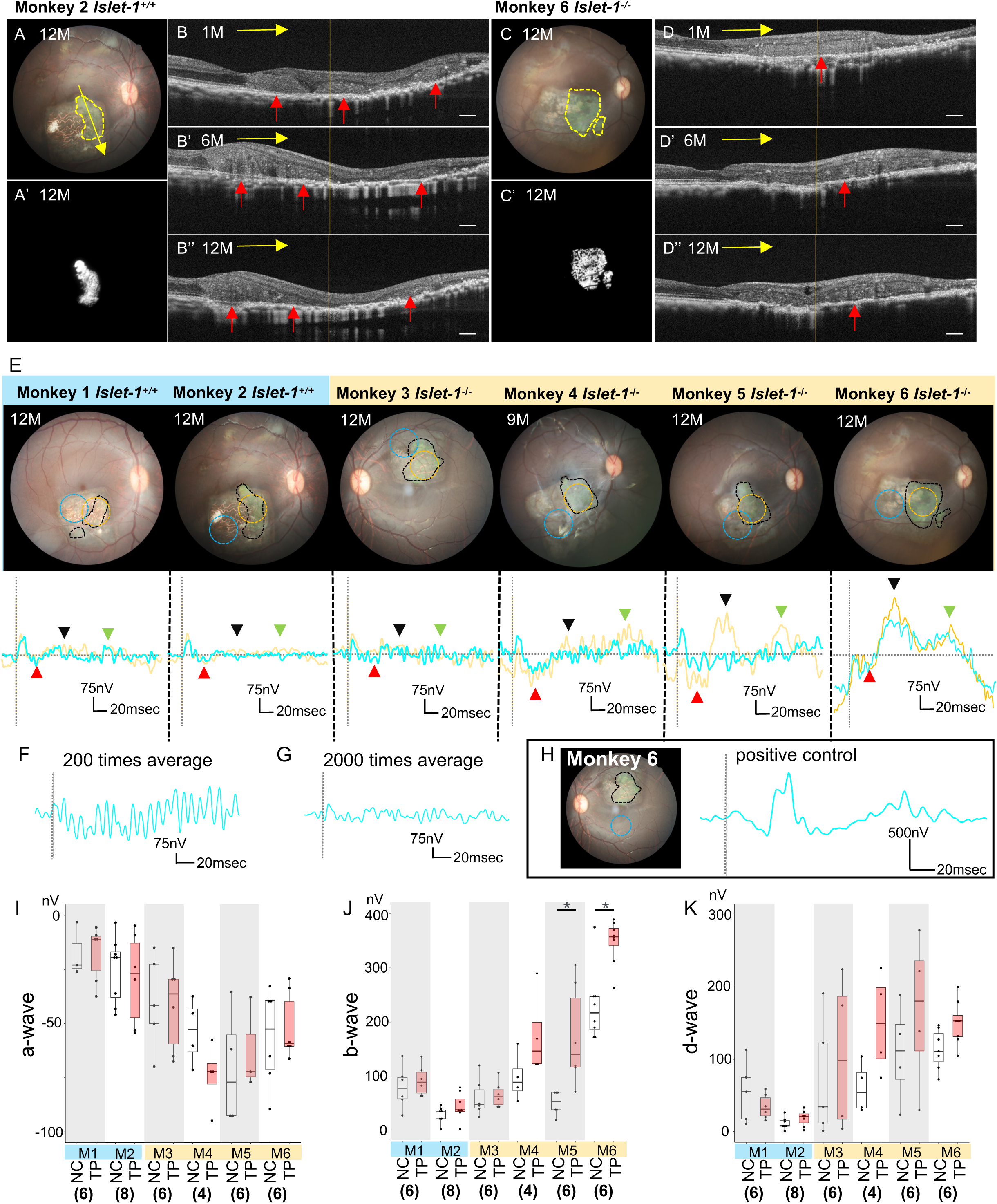
FMERG analysis of transplanted MHP eyes. (A–D’’) Representative imaging of transplanted grafts in monkey eyes. Fundus images (A, C), FA images showing Crx::Venus expression (A’, C’), and OCT images (B–B’’, D–D’’) illustrate the progression of engrafted RO sheets over one year. The yellow dashed line outlines the grafted RO area, while red arrows indicate the graft position post-transplantation. (E) FMERG recordings comparing light responses in grafted (red dot circles) and non-grafted laser-treated (white circles) areas at 6–13 months post-transplantation, using a 10° FMERG field. The black dotted line marks the grafted RO region. Below, representative FMERG waveforms are shown (red arrowhead: a-wave; black arrowhead: b-wave; green arrowhead: d-wave). The blue line indicates the grafted area (TP), while the yellow line represents the non-grafted region (NC). (F–G) Comparison of ERG signal averaging at 200× (F) and 2000× (G) reveals reduced noise with 2000× averaging. (H) Positive control ERG recording with 2000× averaging, obtained from the contralateral retinal area opposite the fovea in the transplanted region. (I–K) Comparison of FMERG wave amplitudes between non-grafted (NC) and grafted (TP) areas across monkeys for a-wave (left), b-wave (middle), and d-wave (right). (*Exact Wilcoxon rank sum test, p < 0.05). Scale bars: 300 μm (B–B’’, D– D’’); 20 μm (E); 25 μm (F–J).

In Monkey 1, half of the graft disappeared between 6 and 8weeks post-transplantation (Supplemental Figure 4A). Although fluorescein angiography (FA) showed no signs of rejection, such as leakage from grafts, OCT revealed increased choroidal thickness at 6 weeks, suggesting potential immune cell accumulation (Supplemental Figure 4B and C). After administering sub-Tenon triamcinolone acetate (STTA) injections, no further graft loss was observed. IHC analysis at one year showed no significant difference in immune cell presence between grafted and intact areas (Supplemental Figure 4D-G).

Following the graft loss in Monkey 1, we refined our protocol by adding periodic local STTA injections alongside systemic cyclosporine administration, maintaining serum levels >70 ng/mL (Supplemental Figure 5A and B). For monkeys with insufficient cyclosporine intake from food, subcutaneous cyclosporine injections were provided based on observed declines in blood levels post-injection (Supplemental Figure 5C and D). In Monkey 3, a combination of oral and subcutaneous cyclosporine from 6 months post-transplantation successfully stabilized blood levels (Supplemental Figure 5E).

Monkey 4 developed postoperative endophthalmitis at 3 weeks post-transplantation, presenting with anterior chamber fibrin masses and vitreous opacity. Treatment included vitrectomy and intravitreal injections of vancomycin and ceftazidime. Although grafts became visible and stabilized, *Crx::Venus* brightness and retinal thickness observed via OCT declined between 7 and 9 months (Supplemental Figure 3K, L, 6A-C). Monkey 4 was euthanized at 9 months post-transplantation, and IHC analysis showed no significant immune cell accumulation, though focal choroidal excavation was noted at the graft site (Supplemental Figure 6C-G). Overall, *Crx::Venus* brightness increased until approximately 13 weeks post-transplantation and remained stable thereafter (Supplemental Figure 3K). Graft thickness similarly increased until 10 weeks and then plateaued (Supplemental Figure 3L).

### Increased b-wave in grafted areas with Islet-1⁻/⁻ RO sheets

We transplanted six RO sheets, covering an area approximately equivalent to the 10-degree recording area for FMERG. FMERG assessments were conducted every two months from 6 to 12 months post-transplantation to evaluate retinal function in the grafted region. We compared FMERG recordings from the grafted and non-grafted areas within the same ONL-ablated lesion in each eye. Initially, signal detection was challenging due to a low signal-to-noise ratio with 200 waveform averaging (Figure 2F). Increasing the averaging to 2000 enabled clear identification of a-, b-, and d-waves, using positive controls with the same averaging for reference (Figure 2G).

The b-wave amplitude was significantly higher in *Islet-1⁻^/^⁻* RO-transplanted areas compared to non-grafted regions, while a-wave and d-wave amplitudes remained unchanged (Figure 2E). Notably, two of four eyes receiving *Islet-1⁻^/^⁻* grafts (Monkey 5 and 6) exhibited b-wave improvements of 21.6% and 24.2%, respectively (Figure 2E and J). In contrast, no FMERG improvements (a-, b-, or d-waves) were observed in *Islet-1⁺^/^⁺* RO-transplanted areas (Figure 2E and I-K). Since *Islet-1⁻^/^⁻* ROs lack ON-BCs and are not expected to generate the b-wave, the observed increase likely originates from host ON-BCs receiving light-evoked electrical responses from grafted photoreceptors.

### Immunohistochemical characterization of Islet-1⁺^/^⁺ and Islet-1⁻^/^⁻ grafts post-transplantation

The eyes of three monkeys, Monkey 1 (*Islet-1⁺^/^⁺),* 3 *(Islet-1^-/-^),* and 4 *(Islet-1^-/-^),* were processed for IHC analysis after euthanasia at 61, 59, and 40 weeks post-transplantation, respectively. Crx::Venus-positive grafted PRCs were dominant in the graft areas of both monkeys (Figure 3A-C’). As expected, PKCα and GNGγ13 (pan-ON BC markers) were present in *Islet-1⁺^/^⁺* grafts but absent in *Islet-1^⁻/⁻^* grafts (Figure 3D-G’’). Notably, PKCα⁺ and GNGγ13⁺ BC dendrites extended into the grafted PRCs, indicating synaptic integration (Figure 3F-G’’, yellow arrows). L/M opsin was expressed in the OS within the graft PRC rosettes, while rhodopsin-positive cells exhibited unpolarized expression, a common feature of stressed PRCs (Figure 4A-C’).

**Figure 3.**
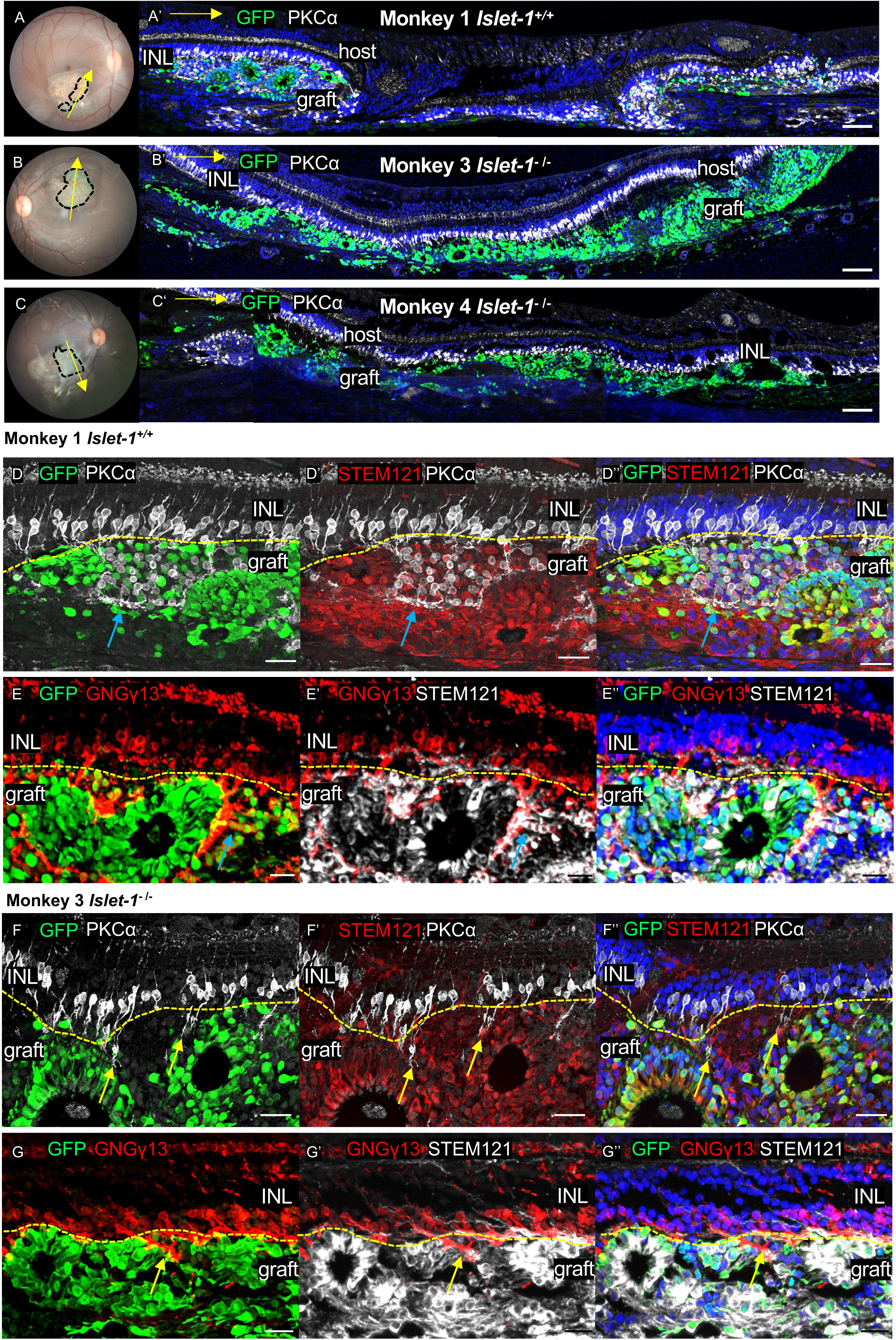
Histological evaluation of human ESC-derived *Islet-1^+/+^* and *Islet1^-/-^* RO sheets after transplantation. (A–C’) Immunostaining of monkey retinas transplanted with *Islet-1^+/+^* (A) or *Islet1^-/-^* (B, C) RO sheets at 12 months (A, B) and 9 months (C) post-transplantation. GFP marks graft-derived PRCs, while PKCα labels BCs. Yellow arrows in the fundus images indicate the orientation of the retinal sections. (D–G’’) Comparison of bipolar cell presence in *Islet-1^+/+^* and *Islet1^-/-^*grafts. PKCα- and GNGγ13-positive BCs (blue arrows) are present in *Islet-1^+/+^*grafts (D– E’’) but absent in *Islet1^-/-^* grafts (F–G’’), confirming the selective depletion of ON-bipolar cells in *Islet1^-/-^*transplants. The dendrite of host BCs extended into the grafted area (yellow arrows). Scale bars: 100 μm (A–C), 30 μm (D–D’’, F–F’’), 20μm (E–E’’, G–G’’).

**Figure 4.**
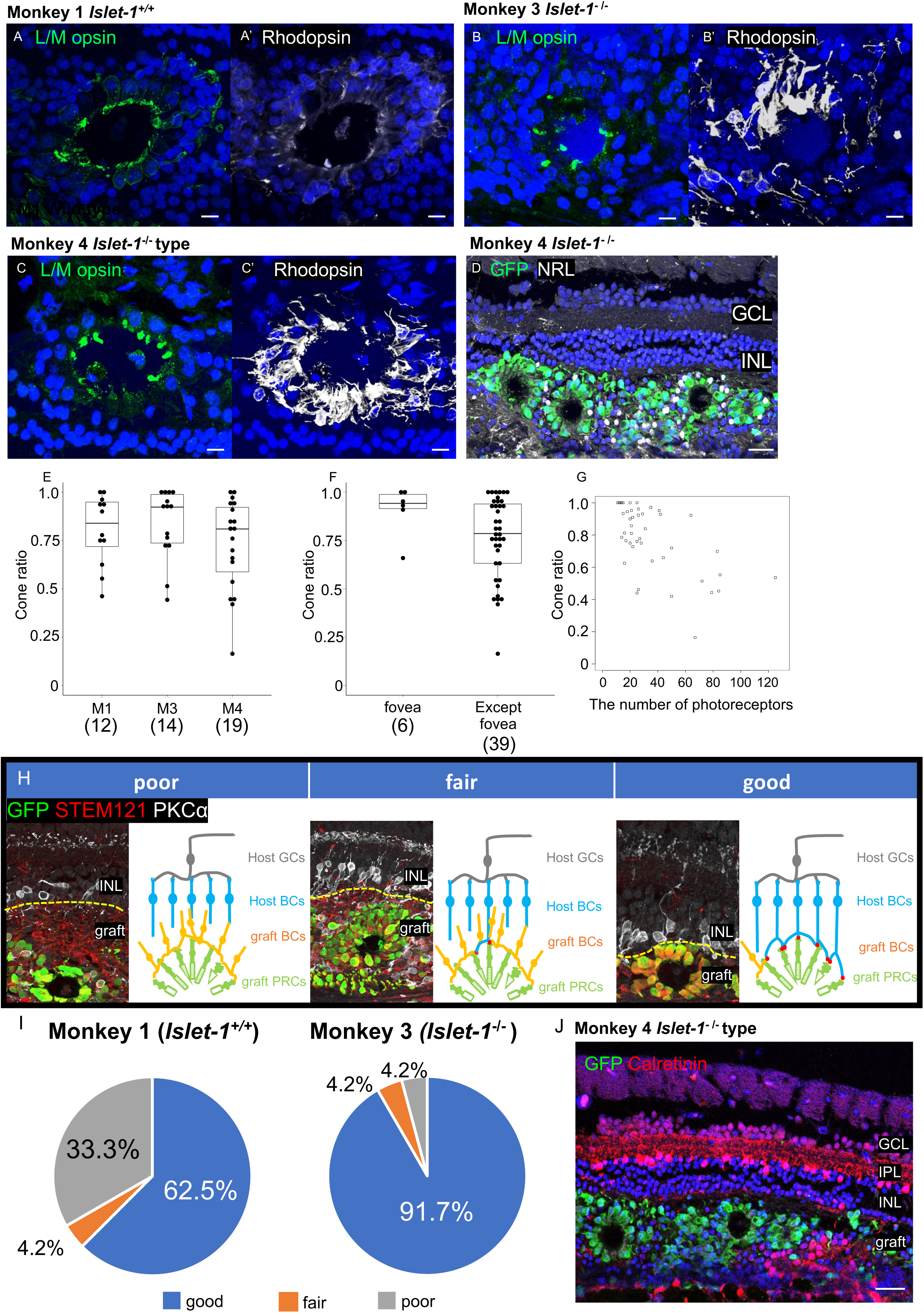
Histological evaluation of cone PRC proportion and host-graft contact. (A–C) Immunostaining of transplanted monkey retinas showing L/M-opsin and rhodopsin expression in *Islet-1^+/+^* and *Islet1^-/-^* grafts post-transplantation, indicating the differentiation of cone and rod PRCs. (D) Immunostaining of GFP and NRL in monkey 4, highlighting graft-derived photoreceptors. (E) Quantification of cone PRC ratio per rosette across individual monkeys (M1: 12 rosettes, M3: 14 rosettes, M4: 19 rosettes). No significant difference was observed (Exact Wilcoxon rank sum test). (F) Comparison of cone PRC ratio per rosette in the fovea versus the parafoveal-perifoveal regions (fovea: 6 rosettes, extra-foveal regions: 39 rosettes). No significant difference was detected (Exact Wilcoxon rank sum test). (G) Correlation analysis between the cone PRC ratio and total PRC count per rosette (n = 45 rosettes, Spearman’s rank correlation coefficient = -0.62), suggesting an inverse relationship between rosette size and cone PRC proportion. (H) Schematic representation of host BC and graft PRC contact patterns. Contact was graded into three categories: Poor (No contact), Fair (Partial contact), and Good (Direct contact). (I) Quantification of host BC-graft PRC contact across monkeys. Monkey 3 exhibited significantly better host-graft synaptic contact than monkey 1 (Islet-1+/+: Good: 62.5%, Fair: 4.2%, Poor: 33.3%; Islet-1-/-: Good: 91.7%, Fair: 4.2%, Poor: 4.2%; n = 24 rosettes per group, p = 0.023, Fisher’s Exact Test for Count Data). (J) Representative immunostaining image of GFP and calretinin-positive cells in monkey 4, illustrating host-graft interaction. Scale bars: 10 μm (A–C’), 30 μm (D), 50 μm (J).

To quantify cone PRCs among Crx::Venus-positive PRCs, we analyzed their spatial distribution. Cone PRCs were primarily located apically, while rod PRCs were positioned basally, mirroring the layered structure of the human retina (Figure 4D). The cone PRC ratio (73%-83%) was much higher than the 10–15% ratio in our previous report (18) (Figure 4E). No significant difference was observed between the foveal (0.91 ± 0.12) and parafoveal- perifoveal (0.76 ± 0.21) regions (Figure 4F). Interestingly, a correlation was observed between PRC rosette size and cone PRC ratio—larger rosettes with more PRCs exhibited a lower cone PRC ratio (Figure 4G). This suggests that rod progenitor cells may have proliferated less efficiently or been lost in smaller rosettes.

### Islet-1⁻^/^⁻ RO transplantation significantly improved contact efficiency between host BCs and graft PRCs

We examined whether *Islet-1⁻^/^⁻*grafts exhibited a higher PRC ratio compared to *Islet-1^⁺/⁺^* grafts, due to the reduced number of BCs within the *Islet-1⁻/⁻* grafts. Indeed, the PRC ratio in *Islet-1⁻^/^⁻* grafts was significantly higher than in *Islet-1^⁺/⁺^* grafts across different monkeys (Supplemental Figure 7A and B). Next, we quantified the extent of direct contact between host BCs and grafted PRC rosettes, categorizing contact efficiency into three grades: good (direct contact), fair (partial contact), and poor (no contact) (Figure 4H). PRCs in *Islet-1⁻^/^⁻* grafts exhibited a significantly higher direct contact ratio with host BCs compared to *Islet-1⁺^/^⁺* grafts (*Islet-1⁺^/^⁺*: Good 62.5%, Fair 4.2%, Poor 33.3%; *Islet-1⁻^/^⁻*: Good 91.7%, Fair 4.2%, Poor 4.2%; *p* = 0.023) (Figure 4I). We further investigated whether graft cell migration disrupted the host IPL, as previously reported in host rat retinas (7). Calretinin immunostaining of the host retina showed an intact IPL structure above the grafts in all the monkey eyes examined (4 sections each from M1, M3, M4) (Figure 4J).

### Multiple BC subtypes extend dendrites to form connections with grafted PRCs in Islet-1⁻^/^⁻ RO

We assessed graft PRC axon elongation and the dendritic extension of host BC subtypes to evaluate host–graft integration with *Islet1****⁻^/^⁻*** transplant. First, we measured graft PRC axon length, defined as the distance from the cytoplasm edge to the axon terminal (Figure 5A and A’), with an average length of 11.9 ± 6.5μm (Figure 4B). Next, we analyzed host BC dendrite extension in the laser-treated area between 9 months and 1 year (9M–1Y) post-treatment.

**Figure 5.**
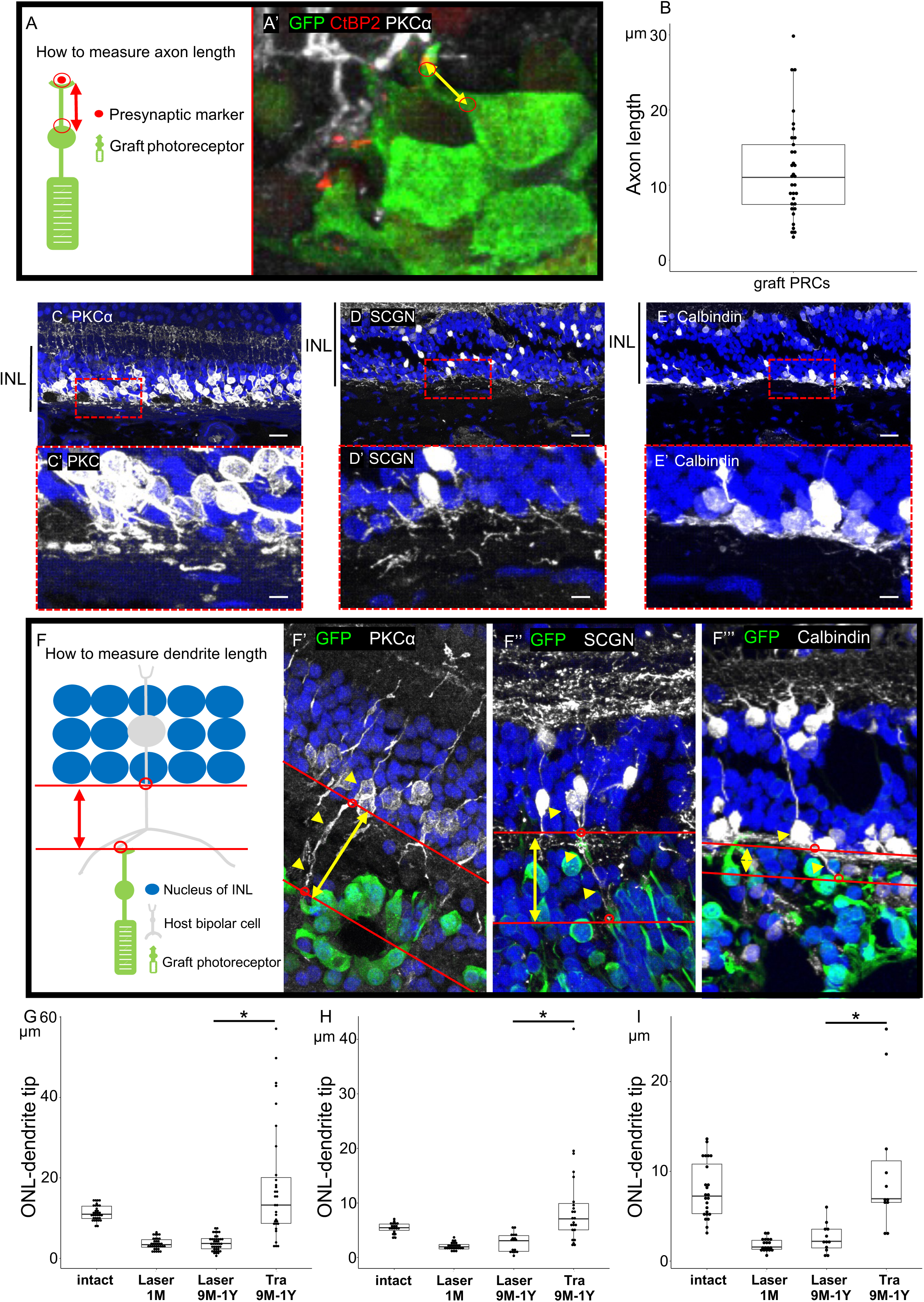
Assessment of graft PRC axon length and host BC dendrite length. (A) Schematic representation illustrating the method for measuring axon length from the axon base to the axon terminal (CtBP2-positive synaptic marker). (B) Quantification of PRC axon length within each graft (n = 32 PRCs, 1 eye). (C–E’) Immunohistochemical analysis of host BC dendrites following 9 months to 1year post-laser treatment. Representative images show dendrites of PKCα-positive BCs (C–C’), SCGN-positive BCs (D–D’), and Calbindin-positive BCs (E–E’). (F) Schematic representation of dendrite length measurement in host BCs. (F’–F’’’) Representative histological images depicting dendrite extension in three BC subtypes (yellow arrowheads). Dendrite length was measured from the bottom of the INL to the axon tips of graft PRCs (yellow double-headed arrows). (G–I) Changes in dendrite length of host BCs before and after laser treatment and post-transplantation. PKCα-positive BCs (Intact: n = 1 eye, 33 BCs; Laser 1M: n = 1 eye, 35 BCs; Laser 9M–1Y: n = 2 eyes, 41 BCs; Transplantation 9M–1Y: n = 1 eye, 31 BCs) in (G). SCGN-positive BCs (Intact: n = 1 eye, 19 BCs; Laser 1M: n = 1 eye, 24 BCs; Laser 9M–1Y: n = 2 eyes, 17 BCs; Transplantation 9M-1Y: n = 2 eyes, 23 BCs) in (H). Calbindin-positive BCs (Intact: n = 1 eye, 24 BCs; Laser 1M: n = 1 eye, 19 BCs; Laser 9M–1Y: n = 2 eyes, 12 BCs; Transplantation 9M–1Y: n = 1 eye, 11 BCs) in (I). Statistical significance was assessed using the Wilcoxon rank sum test (*p <0.05). Scale bars: 20 μm (C, D, E); 5 μm (C’, D’, E’).

Compared to 1-month post-treatment, BC cytoplasmic and dendritic structures appeared disorganized after 9M–1Y (Figure 4C–E’). However, dendrite length in laser-damaged areas without grafts remained unchanged between 1 month and 9M–1Y (Figure 4G-I). In contrast, in grafted areas, the dendrites of three BC subtypes were significantly elongated compared to those in laser-treated but non-grafted areas after 9M–1Y (Figure 4F-I). Interestingly, host BC dendrites in grafted areas were even longer than those in intact regions, suggesting that PKCα-, SCGN-, and Calbindin-positive BCs regrow their dendrites in response to RO engraftment in NHPs.

### Different host BC subtypes contribute to synapse formation with grafted PRCs in Islet-1⁻^/^⁻ RO transplantation

In previous NHP studies, identifying host BC–graft PRC synaptic contacts was challenging due to the presence of graft-derived BCs. However, with *Islet-1⁻/⁻* RO transplantation, where ON-BCs are absent, we could unequivocally observe host PKCα- positive BC dendrites extending toward graft PRCs and forming contacts with CtBP2 (a presynaptic ribbon synapse marker) at the axon terminals of Crx::Venus-positive graft PRCs (Figure 6A–A’’). These extended host PKCα-positive BC dendrites in the graft area were associated with coupling of CtBP2 (presynaptic) and mGluR6 (postsynaptic) markers, indicating synaptic connectivity between host PKCα-positive BCs and graft PRCs (Figure 6B–B’’). Beyond PKCα-positive BCs, additional host BC subtypes, including EAAT2-, Calbindin-, and SCGN-positive BCs, established synaptic connections with Crx::GFP-positive axons (Figure 6C–C’’, Supplemental Figure 8A–B’’) or with graft PRC cone pedicles, often arranged in rosette formations (Supplemental Figure 8C–C’’). Moreover, calbindin-positive HC processes frequently invaginated into Crx::Venus PRC axon terminals, contacting CtBP2-positive synaptic ribbons, suggesting the formation of classic photoreceptor triad synapses involving host BCs, HCs, and graft PRCs (Supplemental Figure 8D-D’’).

**Figure 6.**
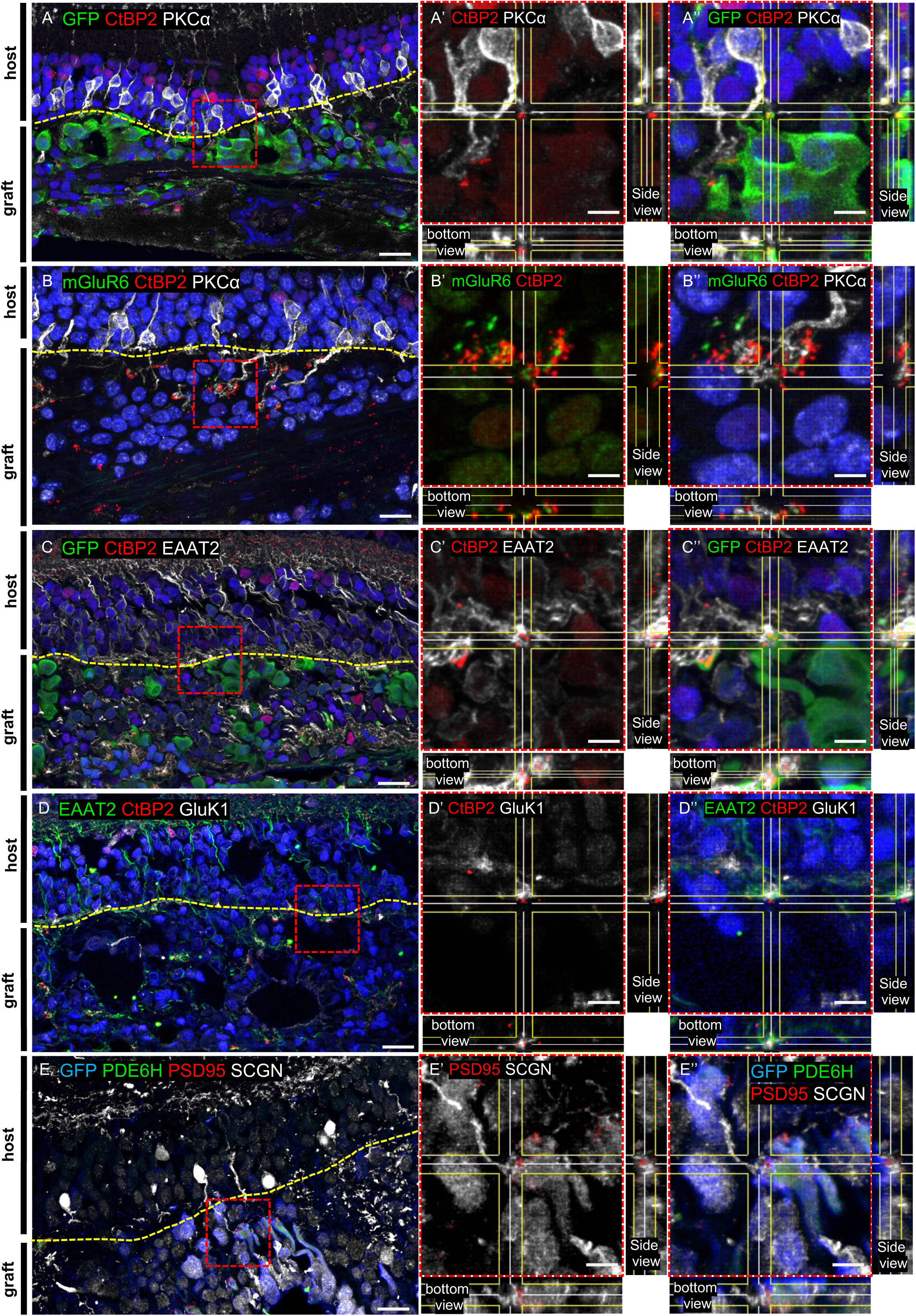
Synaptic connections between host bipolar cells and grafted photoreceptors in *Islet1^-/-^* grafts. (A, C) Presynaptic marker CtBP2 is localized at the margins of GFP-labeled grafted PRCs and the dendritic tips of PKCα-positive bipolar cells (A) and EAAT2-positive BCs (C). (B, D) Co-localization of presynaptic and postsynaptic markers at host BC dendritic tips: CtBP2 (presynaptic) and mGluR6 (postsynaptic) at PKCα-positive BC dendrites (B). CtBP2 (presynaptic) and GluK1 (postsynaptic) at EAAT2-positive BC dendrites (D). (E) Presynaptic marker PSD95 is localized at the margins of GFP- and PDE6H-labeled grafted cone PRCs and at the dendritic tips of SCGN-positive bipolar cells. (A’–A’’, B’–B’’, C’–C’’, D’–D’’, E’– E’’). 3D reconstruction illustrating synaptic contact between grafted PRCs and multiple host BC subtypes. Scale bars: 15 μm (A, B, C, D, E); 4 μm (A’, A’’, B’, B’’, C’, C’’, D’, D’’, E’, E’’).

Furthermore, GluK1, a postsynaptic marker for OFF-BCs, was expressed at the interface between host EAAT2-positive BCs and Crx::Venus-positive PRCs (Figure 6D–D’’). The cone/rod specificity of host BC–graft PRC connections was difficult to determine due to overlapping BC marker expression in both cone and rod BCs. Nevertheless, we observed that cone-specific SCGN-positive BCs formed synaptic contacts with graft-derived cone PRCs, identified by PDE6H and GFP expression, suggesting de novo reconstruction of cone circuitry (Figure 6E–E’’).

### Electron microscopy confirms synaptic integration between host BCs and grafted PRCs

To further validate host BC–graft PRC synapse formation, we performed electron microscopy (EM) analysis at 55 weeks post-transplantation, guided by Crx::Venus fluorescence and DAPI staining prior to embedding (Figure 7A and B, Supplemental Figure 9A). Using serial EM image reconstruction, we traced graft-derived PRCs and their axons extending toward the host INL (Figure 7C and D, Supplemental Figure 9B). Notably, synaptic ribbons frequently formed multiple triad synapses within graft-derived PRC pedicles, where BCs and HCs made putative invaginating contacts (Figure 7E and F, Supplemental Figure 9C–F). Since ON-BCs were absent in *Islet-1⁻/⁻* grafts (Figure 3F–G’’), these invaginating BC processes were likely from host ON-BCs. Additionally, we employed Serial Block-Face Scanning Electron Microscopy (SBFSEM) to examine graft PRC–host BC connectivity. This high-resolution imaging method allows precise neuron tracing to evaluate synaptic interactions. Imaging the direct contact area between graft PRCs and host BCs (Figure 7G and H), we identified host ON-BCs and OFF-BC making synaptic connections with transplanted cone PRC axon terminals (Figure 7I). Importantly, the dendrites of ON-BC formed invaginating synapses with the transplanted cone cell, while the dendrites of OFF-BC made flat contact with the transplanted cone cell. (Figure 7J and K).

**Figure 7.**
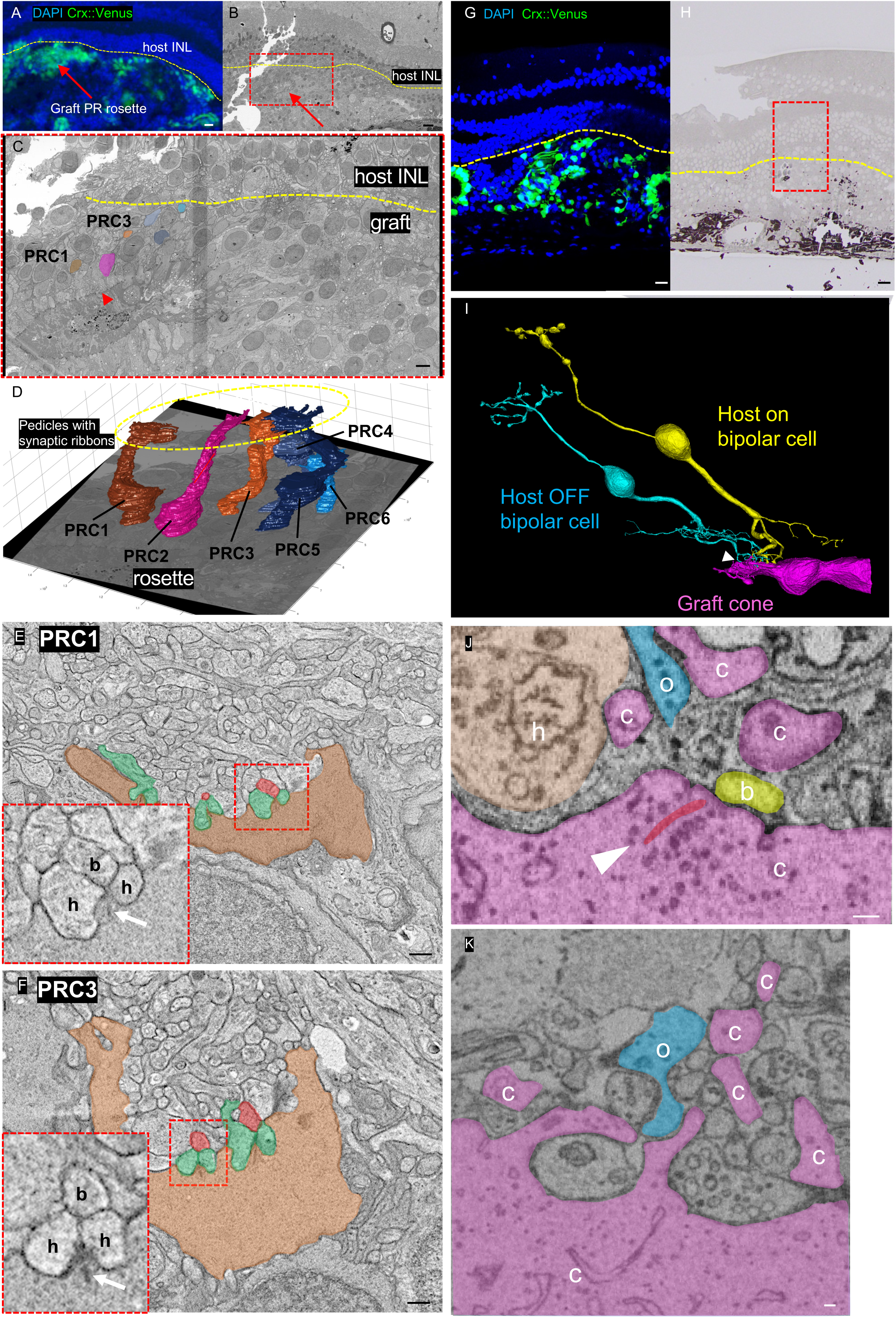
Synaptic Microstructures between hot BCs and grafted PRCs. (A) Pre-embedding images labeled with DAPI and Crx::Venus, showing the rosette structure (red arrow) used for tracing grafted PRCs. (B) EM image of the same area shown in (A), with the rosette (red arrow) corresponding to the one identified in (A). (C) Enlarged view of the region within the red dotted rectangle in (B). Red arrowhead showed external limiting membrane of rosette. (D) 3D reconstruction showing graft PRCs extending from the rosette into the host INL using tomography; six PRCs were traced. (E, F) EM images of individual PRC axon terminals: PRC1 (brown, E) and PRC3 (orange, F). The red dotted rectangle highlights a triad synapse, where bipolar cells (b) and horizontal cells (h) invaginate into the ribbon synapse (white arrow). (G) Pre-embedding images labeled with DAPI and Crx::Venus. (H) EM image of the corresponding region in (G). (I) 3D reconstruction from SBFSEM analysis, revealing direct contact (white arrowhead) between a host ON bipolar cell, a host OFF bipolar cell, and a grafted cone PRC. (J) Higher magnification image highlighting the contact site between the grafted cone PRC (pink) and the host ON bipolar cell (yellow). The synaptic ribbon (white arrowhead, red coloring) is positioned opposite the host ON bipolar cell process, confirming synaptic integration. (K) Higher magnification image showing the flat contact between the grafted cone PRC (pink) and the host OFF bipolar cell (blue). c, cone cell; b, On bipolar cell; o, Off bipolar cell; h, horizontal cell. Scale bars: 20 μm (A, B, G, H); 5 μm (C); 500 nm (E, F); 50 nm (J, K).

## Discussion

In this study, we provide the first demonstration that host BCs in the primate macula can regenerate dendrites and form functional synapses with transplanted human photoreceptors. By using genome-edited *Islet-1⁻/⁻* ROs that lack ON-BCs, we eliminated intra-graft competition and directly visualized host-driven synaptogenesis. Recovery of focal macular ERG b-waves, together with ultrastructural evidence of triad synapses and ON- and OFF host-graft rewiring, establishes that transplanted organoids can reconstitute cone pathways in the primate central retina. These findings close a critical knowledge gap, showing that cone BC plasticity— previously uncertain in primates—can be harnessed for vision restoration.

Functional recovery was modest but significant. Two of four eyes receiving *Islet-1⁻/⁻* grafts exhibited measurable increases in b-wave amplitude, whereas *Islet-1⁺/⁺* grafts failed to present detectable recovery. It is important to note that in-vivo electrophysiological recordings may underestimate the graft efficacy, as they tend to be less sensitive compared to subjective examinations (24). In rodents, we have consistently shown the recovery of host RGC responses in the grafted area using the multiple electrode arrays (6–8), despite the difficulty in recording these responses in vivo. Because *Islet-1⁻/⁻* organoids lack ON-BCs, these responses most likely reflect host cone bipolar activity driven by grafted photoreceptors. The absence of a-wave improvement may relate to the rosette-like architecture of transplanted organoids, which disrupts photoreceptor orientation and outer segment maturation. These observations highlight the b-wave as the most reliable electrophysiological measure of host–graft connectivity in the primate macula.

At the structural level, multiple host BC subtypes (PKCα⁺ rod BCs, SCGN⁺ and EAAT2⁺ cone BCs, and Calbindin⁺ BCs) extended dendrites into grafts and formed synaptic contacts with graft derived RPC terminals. Electron microscopy studies confirmed both invaginating ON-BC and flat OFF-BC synapses with graft-derived cones, demonstrating reconstitution of canonical photoreceptor–bipolar–horizontal cell triads. These findings indicate that even foveal midget BCs, once thought to have limited rewiring capacity (16), retain regenerative potential when stimulated by developing grafts. Interestingly, grafts contained an unexpectedly high proportion of cones (∼80%) compared to that in our previous report (20), which may reflect the influence of the foveal microenvironment or differential vulnerability of rods under inflammatory stress. Regardless of the mechanism, cone enrichment is favorable for restoring central vision.

Immunological management posed another challenge. Although most grafts survived long-term under systemic and local immunosuppression, persistent macular edema and one case of graft loss suggested localized immune activation in the foveal region. This finding aligns with our previous report, which indicated a possible immune reaction following retinal organoid sheet transplantation at the fovea in the NHP macular hole model (25). This contrasts with our prior peripheral xenotransplantation studies (20, 24), which showed minimal immune reaction, and suggests that the macula may be more immunologically vulnerable. Refined surgical approaches and optimized immunomodulation will likely be required. Importantly, however, our recent clinical trial of allogeneic retinal organoid transplantation near the fovea showed no rejection (9), and prior fetal retinal transplant studies similarly reported low rejection rates (26, 27). Furthermore, ESC/iPSC-derived organoids exhibit low immunogenicity, with reduced HLA expression and secretion of immunosuppressive factors (28, 29). These observations support the feasibility of clinical translation, though surgical trauma near the insertion site may still trigger immune responses. Avoiding graft placement adjacent to entry sites may mitigate this risk (30).

A limitation of this study is the use of an acute laser-ablation model, which differs from the progressive remodeling and gliotic sealing that occur in hereditary retinal degeneration (31, 32). Nonetheless, the model recapitulated essential features, e.g., BC dendrite retraction and postsynaptic protein loss, and allowed us to test whether developing ROs could stimulate host plasticity. Human retinitis pigmentosa typically progresses over decades and shows regional variation within the same eye, including gliotic barriers that may impede integration (31). In our previous clinical trial, we placed the RO grafts over areas of relatively preserved retinal pigment epithelium, which achieved stable engraftment by forming an outer plexiform layer-like structure between the host and graft tissues with no scarring barriers (9, 32) similar to the outcomes observed in animal studies. This suggests that the transplantation site and disease stage are critical determinants of success. Future studies in chronic genetic NHP models will be needed to confirm whether the rewiring observed here is durable in long-standing degeneration.

In conclusion, these findings provide proof-of-concept that primate cone and rod BCs can re-establish functional synapses with transplanted human PRCs. The enhanced integration observed with *Islet-1⁻/⁻* grafts suggests that targeted genetic modification of donor ROs may further optimize connectivity. While additional studies in chronic models are needed, the demonstration of functional synaptic integration in the primate macula represents a pivotal step toward clinical application of genome-edited retinal organoid transplantation, particularly for patients with macular diseases in which cone-mediated central vision is lost.

## Methods

### Sex as a biological variable

Our study used Male monkeys to ensure consistency across experimental conditions by reducing sex-related variability.

### Human ESC line

Human ESC (KhES-1) line was established at and distributed by Kyoto University (NIH Approval Number: NIHhESC-22-0487). The KhES-1 Crx::Venus line (RIKEN BioResource Center, Cell Number: HES0653) (33) was used to generate the genome-edited *Islet-1*^-/-^ line in our previous study (7). KhES-1 Crx::Venus line and *Islet-1*^-/-^ line were used in this study in accordance with human ESC research guidelines of the Japanese government.

### Animals

A total of 7 cynomolgus monkeys (provided by Eve Bioscience) were used in the experiment (one for laser treatment, and Monkey 1 to Monkey 6 for transplantation; Supplemental Table 1). Each monkey was transplanted with 6 RO sheets; Monkey 1 and Monkey 2 were transplanted with *Islet-1^+/+^* retinal organoid sheets. Monkey 3 to Monkey 6 were transplanted with *Islet-1*^-/-^ retinal organoid sheets.

### Cell culture and differentiation of human ESCs

Retinal differentiation was performed using the modified SFEBq method (34). In brief, subconfluent human ESCs were cultured with 5 μM SB431542 (Sigma-Aldrich, catalog S4317) and 300 nM SAG (Enzo Biochem, catalog ALX-270-426-M001) from 24 h prior to differentiation. Cells were then dissociated into single cells using TrypLE Select Enzyme (Thermo Fisher Scientific, catalog A12859-01), suspended in 100 μL growth factor-free chemically defined medium (gfCDM; Ham’s F-12 Nutrient Mix (Life technologies, catalog 11765-054); IMDM+ GlutaMAX (Life technologies, catalog 31980-030); CD lipid concentrate (Life technologies, catalog 11905-031); Knockout Serum Replacement (Life technologies, catalog 10828-028); 1-thioglycerol (Sigma, catalog M6145)) with 10 μM Y-27632 (FUJIFILM Wako, catalog 253-00513), and seeded in low cell adhesion 96-well V-bottomed plates (Sumitomo Bakelite, catalog MS-9096V) at 1.2×10^4^/well. On differentiation day (DD) 3, aggregates were treated with 1.5 nM recombinant human BMP4 protein (R&D Systems, catalog HZ-1045). The gfCDM medium was half-changed every 3 days from DD6 to DD15. Aggregates on DD18 were transferred to DMEM/F12+Glutamax (Invitrogen, catalog 10565-042) with 3μM CHIR99021 (FUJIFILM Wako, catalog 034-23103), 5 μM SU5402 (FUJIFILM Wako, catalog 193-16733), and N2- supplement (100×, Life technologies, catalog 17502-048) for 3–4 days in a 90-mm low adhesion culture dish (Corning International). Subsequently, aggregates were cultured in the control maturation medium (DMEM/F12 with GlutaMAX (Life technologies, catalog 10565-042); N2 supplement (Life technologies, catalog 17502-048); 10%FBS (Gibco, catalog 10270-106), 100μM Taurin (Sigma, catalog T8691-25G), Penicilin-Streptomycin-AmphotericinB Suspension (FUJIFILM Wako, catalog 161-23181. The medium was exchanged every 3-4days until use at approximately DD60.

### Laser induced retinal degeneration model

To establish a laser-induced retinal degeneration model, we employed the 577-nm OPSL laser system (PASCAL Streamline yellow, Topcon) for selective photocoagulation of the ONL using a laser contact lens (macula lens 901, HAAG-STREIT INTERNATIONAL) with a magnification of 0.96×. The pattern scan consisted of 9 spots arranged in a 3 × 3 grid. The parameters for the laser procedure included a spot spacing of 0μm, a laser size of 100μm, and a duration of 15ms. We evaluated the photocoagulated area immediately after the laser with Spectral Domain-OCT (SD-OCT) (RS3000, NIDEK) and fundus camera (CX-1, Canon), and we added the laser if deemed necessary.

### Procedure of transplantation

The surgery was performed following the previous clinical study by Hirami et al (9). A localized retinal detachment was created in the laser area by subretinal injection of oxiglutatione intraocular irrigating solution (BSS Plus, Alcon) using a 38GPolyTip Cannula (MedOne) The cannula hole was enlarged by scissors (Pinnacle 360^TM^ 25G Vertical Scissors, Synergetics) for graft insertion using a customized 24G plastic cannula. Another hole was created within the retinal detachment to allow smooth graft insertion without increasing the subretinal pressure. Grafts were transplanted into the subretinal space using a 1:6 sodium hyaluronate and chondroitin sulfate sodium (Viscoat, Alcon): BSS Plus, then spread throughout the target area with subretinal forceps (keeler). Perfluoro-n-octane (perfluoron, Alcon) was injected to drain subretinal fluid, followed by silicon oil (Alcon) injection after fluid-air exchange.

### Immunosuppression

Immunosuppression was achieved through sub-Tenon triamcinolone acetonide (STTA) injections (KENACORT-A 40mg/mL, 0.5ml, Bristol Myers Squibb) administered at the end of transplantation surgery two weeks post transplantation, and at the time of silicon oil removal. STTA injections were also utilized when transplant rejection was suspected, with subsequent administration every two months. Cyclosporine (Sandimmun, NOVARTIS, 10mg/mL, 7-9mL) was administered orally as systemic immunosuppression to maintain serum levels between 70-1000 ng/mL. Subcutaneous cyclosporine injection (10mg/ml 0.5mg/mL) was used when oral administration was difficult to manage. The immunosuppression management protocol is shown in Supplemental Figure 5.

### FMERG recordings

To verify the coagulation of photoreceptors induced by laser treatment, focal macular electroretinography (FMERG) was performed using an ER-80 system (KOWA). FMERG was recorded using a Burian-Allen bipolar contact lens electrode (HANSEN Ophthalmic Development Laboratory) with the ground electrode placed on the ear. Recordings were obatained from the photocoagulated area using the following parameters; stimulus light intensity (39cd/m^2^), background light intensity (1,28cd/m^2^), stimulus duration (100ms) and a 10°stimulus spot. For creating the laser-induced retinal degeneration model, the stimulus repetition rate was 5Hz, and 200 responses were averaged for each recording using a Neuropack MEB-2200 system (Nihon Kohden) For assessments conducted 6 months after transplantation, the stimulus repetition rate was 4.5Hz, and 2000 responses were averaged using a PuREC system (Mayo).

### Immunohistochemistry

The eyes were immediately harvested upon euthanization, fixed in SUPERFIX (Kurabo) for 1 week, and transferred in the phosphate-buffered saline (PBS) without calcium and magnesium. Eyes were embedded in paraffin and sectioned at 10μm thickness using an automated slide preparation system (AS-200; Kurabo). After deparaffinization, 4 types of antigen retrieval methods were employed depending on the antibody used: (1) microwave treatment for 20 minutes in10mM Citrate buffer (pH6.0) at 100℃: (2) autoclave treatment for 20 minutes in10mM Citrate buffer (pH6.0) at120℃ (3) autoclave treatment for 20 minutes in 10mM Trypsin-EDTA (pH9.0) at 120℃:(4) Enzymatic treatment for 10-15 minutes using10μg/mL ProK in 50mM Tris buffer (pH8.0) at 37℃.

Specimens were blocked with blocking solution (1%BSA in PBS-0.3%Tx) for 1hour at room temperature (RT) followed by the incubation with primary antibodies (Supplemental Table 2) over night at 4℃, then with secondary antibodies for 1 hour at room temperature. The images were acquired using LSM 700 (Zeiss) and TCS SP8 (Leica) confocal microscopy platforms. Primary antibodies are listed in Table S2.

### Electron Microscopy

The cornea and lens were removed from the transplanted eye of Monkey 5 and fixed in 2% glutaraldehyde in 0.1M cacodylate buffer at room temperature for three hours. The transplantation area was trimmed and embedded in agarose gel to prepare 300μm sections using a vibratome (DSK-Pro7, Dosaka EM), and stained with Dapi for observation under fluorescence microscope to confirm Crx::Venus positive graft integration in the host retina. The sections were imaged using TCS SP8 (Leica) confocal microscopy and then stored in 4% glutaraldehyde in 0.1 M sodium cacodylate buffer until use. The sections were washed five times with 0.1M cacodylate buffer, followed by fixation in 1% osmium tetroxide (OsO4) in 0.1M cacodylate buffer at 4℃ for one hour and subsequent washing five times with distilled water. The specimens were incubated in 1% uranyl acetate overnight, then washed five more times with distilled water. We dehydrated the specimens in a graded ethanol series (20%, 50%, 70%, 90%, 99.5%, and 100%, each for 5 minutes on ice), followed by 100% acetone for 10min at room temperature. Infiltration with Epon812 (TAAB) ensued, and polymerization was performed at 60℃ for 72 hours. Samples were sectioned at 250nm thickness using ultramicrotome (ULTRACUT-UCT, Leica) equipped with a diamond knife (Diatome), stained with uranyl acetate and lead nitrate, and observed under a field-emission scanning electron microscope (MERLIN VP compact, Carl Zeiss) at 5kV using a backscattered electron detector.

### Serial Block-face Scanning Electron Microscopy (SBFSEM)

One of the vibratome sections from above was shipped to National Institute for Physiological Sciences, Okazaki in a temperature-controlled container for SBFSEM. Retinal pieces were incubated in 1.5% potassium ferrocyanide and 2% OsO4 in phosphate buffered saline for 1 h. After washing, the tissue was placed in a freshly made thiocarbohydrazide solution (0.1 g TCH in 10 ml dH2O heated to 65 °C for 1 h) for 20 min at room temperature. After rinsing at RT, the tissue was incubated in 2% OsO4 at RT for 30 min, rinsed again, and stained en bloc in 2%uranyl acetate overnight at 4℃. After washing, the tissue was stained with Walton’s lead aspartate (0.66% lead in 0.03M L-aspartic acid, pH 5.0∼5.5) at 50℃ for 2 h. After a final wash, the retinal pieces were dehydrated in a graded ice-cold alcohol series and finally embedded in Durcupan resin. Light microscopic images were acquired from semi-thin sections of the resin-embedded tissue surface, and the resin block was trimmed to remain the retinal region overlapped with fluorescent signals of grafted GFP-positive cells. The SBFSEM observation of the retinal region was performed using a Merlin scanning electron microscope (Carl Zeiss) equipped with a 3View in-chamber ultramicrotome system (Gatan). Serial electron microscopic images at 60nm thickness were acquired, and the images were composed of a montage of 4 x 7 tiles, each of which were sized 6144 x 6144 pixels at 5nm / pixel resolution. Image stacks were stitched, concatenated and aligned using TrackEM2 (ImageJ/Fiji). Retinal cells were manually segmented and reconstructed using TrakEM2.

### Statistics

The length of BCs dendrites was quantified from Z-stack images of sectioned retinas from monkey for laser treatment (Figure 1) and monkey 3 and 4 (Figure 4). Orthogonal sections intersecting grafts were resliced in Image J, and the dendrites of BCs were traced manually. BCs were found in three to four separate sections, and the length from the base of the INL to the dendrite tip was measured for each BC.

PRCs axon length was quantified from Z-stack images of sectioned retinal from monkey 4 (Figure 4). Orthogonal sections intersecting grafts were resliced in Image J and PRC axons were traced manually. PRC axons were identified in four separate sections, and the length from the axon base to tip was measured for each PRC.

Graft thickness was measured from the same area in each monkey using OCT. (Supplemental Figure 3L) The longest graft length was measured in each monkey and OCT images were quantified using Image J. Fluorescein angiography (FA) images of grafts were captured using CX-1. (Supplemental Figure 3K), and the mean brightness of Crx::Venus expression in grafts was quantified using Image J.

Cone ratio was measured by GFP and NRL staining. (Supplemental Figure 7D-G) GFP was expressed in graft PRCs, while NRL was expressed in rod PRCs. The cone ratio was calculated as (GFP-positive PRCs-NRL-positive PRCs/ GFP-positive PRCs). Four sections from separate areas from each monkey were stained and all rosettes were manually counted in sectioned retina using Image J.

PRCs ratio was measured using GFP and STEM121 staining. (Supplemental Figure 8A and B) GFP was expressed in graft PRCs, while STEM121 was expressed in all graft cells. The PRCs ratio was calculated as (GFP-positive PRCs/ STEM121-positive graft cells). Between 9-18 sections from separate areas of each monkey were stained. Blood vessels were removed after converting image to 8 bit format. GFP and STEM121 were binarized, and the areas were measured using Image J.

Synaptic contact was evaluated using GFP, STEM121, and PKC staining. (Supplemental Figure 8C and D) Host and graft BCs were distinguished using STEM121 staining. Between 11-12 sections from separate areas of monkey 1 and 4 were stained. Two investigators independently evaluated synaptic contact using a three-point grading scale.

Statistical analyses were conducted using R (version 4.1.2, https://cran.r-project.org/bin/macosx/). The exact Wilcoxon rank-sum test was used for most comparisons, while Fisher’s exact test was used for synaptic contact analysis. Statistical significance was defined as *P* ≤ 0.05.

### Study approval

All animal experiments were conducted in accordance with local guidelines and the ARVO Statement on the Use of Animals in Ophthalmic and Vision Research. All of the experimental protocols were approved by the Institutional Animal Care and Use Committee of RIKEN Kobe Branch.

## Data availability

The datasets supporting the current study have not been deposited in a public repository due to their large size but are available from the corresponding author upon reasonable request. Any additional information required to reanalyze the data is also available upon request.

## Author contributions

Conceptualization, A.O.,Y.F. and M.M.; Data curation, A.O., R.A.,S.Y., and M.M.; Formal Analysis, A.O., A.K., R.A., and M.M; Funding acquisition, Y.F., R.A. N. O. and M.M.; Investigation, A.O., K.K., A.K.,R.A., D.P., N.O.,S.O., S.Y.,S.I., Y.K., and M.M; Methodology, R.A., N.O.,T.M.,S.Y., M.K., Y.K. and M.M.; Project administration, M.M.; Resources, M.K., T.M. S.Y., M.K., and M.M.; Supervision, M.M.; Visualization, R.A. and A.O.; Writing – original draft, A.O and R.A.; Writing – review and editing, Y.F, and M.M.

## Acknowledgements

Animal experiments were conducted at RIKEN BDR with the support of Hiroshi Kiyonari and Masayo Takahashi. We thank Satoshi Shirae and Toshika Senba at Vision Care Co. for their professional support in animal experiments, Michiru Matsumura and Mitsunari Nishida at Kobe City Eye Hospital for cell culture of human ESC and retinal organoid, and Atsuko Imai and Nobuko Hattori at National Institute for Physiological Sciences for SBFSEM analysis.

This research was funded by National Institutes of Health (NIH) grant 1U24EY033272 (Y.F. and M.M.), KAKENHI grant number 23K15925 (R.A.), and Beyer Retina Award (R.A.). This work was partly supported by JSPS KAKENHI Grant Number JP22H04926 (N. O), Grant-in-Aid for Transformative Research Areas ― Platforms for Advanced Technologies and Research Resources “Advanced Bioimaging Support” and Cooperative Study Programs of National Institute for Physiological Sciences (R.A).

## Conflict of Interest

The authors have declared that no conflict of interest exists.

## Notes

### Competing Interest Statement

The authors have declared no competing interest.

## References

1. Marc RE, et al. Neural remodeling in retinal degeneration. Prog Retin Eye Res. 2003;22(5):607–55.

2. Pfeiffer RL, et al. Persistent remodeling and neurodegeneration in late-stage retinal degeneration. Prog Retin Eye Res. 2020;74:100771.

3. Mandai M, et al. iPSC-Derived Retina Transplants Improve Vision in rd1 End-Stage Retinal-Degeneration Mice. Stem Cell Reports. 2017;8(1):69–83.

4. Mandai M. Pluripotent stem cell-derived retinal organoid/cells for retinal regeneration therapies: A review. Regen Ther. 2023;22:59–67.

5. Ozaki A, et al. hPSC-based treatment of retinal diseases - Current progress and challenges. Adv Drug Deliv Rev. 2025;221:115587.

6. Matsuyama T, et al. Genetically engineered stem cell-derived retinal grafts for improved retinal reconstruction after transplantation. iScience. 2021;24(8):102866.

7. Yamasaki S, et al. A Genetic modification that reduces ON-bipolar cells in hESC-derived retinas enhances functional integration after transplantation. iScience. 2021;25(1):103657.

8. Watanabe M, et al. Transplantation of genome-edited retinal organoids restores some fundamental physiological functions coordinated with severely degenerated host retinas. Stem Cell Reports. 2025;20(2):102393.

9. Hirami Y, et al. Safety and stable survival of stem-cell-derived retinal organoid for 2 years in patients with retinitis pigmentosa. Cell Stem Cell. 2023;30(12):1585–1596.e6.

10. Beier C, et al. Deafferented Adult Rod Bipolar Cells Create New Synapses with Photoreceptors to Restore Vision. J Neurosci. 2017;37(17):4635–4644.

11. Shen N, et al. Kerschensteiner D. Homeostatic Plasticity Shapes the Retinal Response to Photoreceptor Degeneration. Curr Biol. 2020;30(10):1916–1926.e3.

12. Fitzpatrick MJ, Kerschensteiner D. Homeostatic plasticity in the retina. Prog Retin Eye Res. 2023;94:101131.

13. Sullivan RK, et al. Dendritic and synaptic plasticity of neurons in the human age-related macular degeneration retina. Invest Ophthalmol Vis Sci. 2007;48(6):2782–91.

14. Sethi CS, et al. Glial remodeling and neural plasticity in human retinal detachment with proliferative vitreoretinopathy. Invest Ophthalmol Vis Sci. 2005;46(1):329–42.

15. Eliasieh K, et al. Cellular reorganization in the human retina during normal aging. Invest Ophthalmol Vis Sci. 2007;48(6):2824–30.

16. Akiba R, et al. Cellular and circuit remodeling of the primate foveal midget pathwayafter acute photoreceptor loss. Proc Natl Acad Sci U S A. 2024;121(37):e2413104121.

17. Akiba R, et al. Host-Graft Synapses Form Functional Microstructures and Shape the Host Light Responses After Stem Cell-Derived Retinal Sheet Transplantation. Invest Ophthalmol Vis Sci. 2024;65(12):8.

18. Watanabe M, et al. Graft-derived horizontal cells contribute to host-graft synapses in degenerated retinas after retinal organoid transplantation. Stem Cell Reports. 2025;20(7):102545.

19. Bringmann A, et al. The primate fovea: Structure, function and development. Prog Retin Eye Res. 2018;66:49–84.

20. Shirai H, et al. Transplantation of human embryonic stem cell-derived retinal tissue in two primate models of retinal degeneration. Proc Natl Acad Sci U S A. 2016 ;113(1):E81–90.

21. Care RA, et al. Partial Cone Loss Triggers Synapse-Specific Remodeling and Spatial Receptive Field Rearrangements in a Mature Retinal Circuit. Cell Rep. 2019;27(7):2171–2183.e5.

22. Dunn FA. Photoreceptor ablation initiates the immediate loss of glutamate receptors in postsynaptic bipolar cells in retina. J Neurosci. 2015;35(6):2423–31.

23. Kalloniatis M, et al. Using the rd1 mouse to understand functional and anatomical retinal remodelling and treatment implications in retinitis pigmentosa: A review. Exp Eye Res. 2016;150:106–21.

24. Tu HY, et al. Medium- to long-term survival and functional examination of human iPSC- derived retinas in rat and primate models of retinal degeneration. EBioMedicine. 2019;39:562–574.

25. Iwama Y, et al. Transplantation of human pluripotent stem cell-derived retinal sheet in a primate model of macular hole. Stem Cell Reports. 2024;19(11):1524–1533.

26. Radtke ND, et al. Vision improvement in retinal degeneration patients by implantation of retina together with retinal pigment epithelium. Am J Ophthalmol. 2008;146(2):172–182.

27. Das T, et al. The transplantation of human fetal neuroretinal cells in advanced retinitis pigmentosa patients: results of a long-term safety study. Exp Neurol. 1999;157(1):58–68.

28. Yamasaki S, et al. Low Immunogenicity and Immunosuppressive Properties of Human ESC- and iPSC-Derived Retinas. Stem Cell Reports. 2021;16(4):851–867.

29. Uyama H, et al. Competency of iPSC-derived retinas in MHC-mismatched transplantation in non-human primates. Stem Cell Reports. 2022;17(11):2392–2408.

30. Petrash CC, et al. Immunologic Rejection of Transplanted Retinal Pigmented Epithelium: Mechanisms and Strategies for Prevention. Front Immunol. 2021;12:621007.

31. Fariss RN, et al. Abnormalities in rod photoreceptors, amacrine cells, and horizontal cells in human retinas with retinitis pigmentosa. Am J Ophthalmol. 2000;129(2):215–23.

32. Ishikura M, et al. Adaptive Optics Optical Coherence Tomography Analysis of Induced Pluripotent Stem Cell-Derived Retinal Organoid Transplantation in Retinitis Pigmentosa. Cureus. 2024;16(7):e64962.

33. Nakano T, et al. Self-formation of optic cups and storable stratified neural retina from human ESCs. Cell Stem Cell. 2012;10(6):771–785.

34. Kuwahara A, et al. Generation of a ciliary margin-like stem cell niche from self-organizing human retinal tissue. Nat Commun. 2015;6:6286.

