## Supplementary figures and tables for "Genome-edited retinal organoids restore host bipolar connectivity in the primate macula"

Supplementary Figure 1–9
Supplementary Table 1, 2

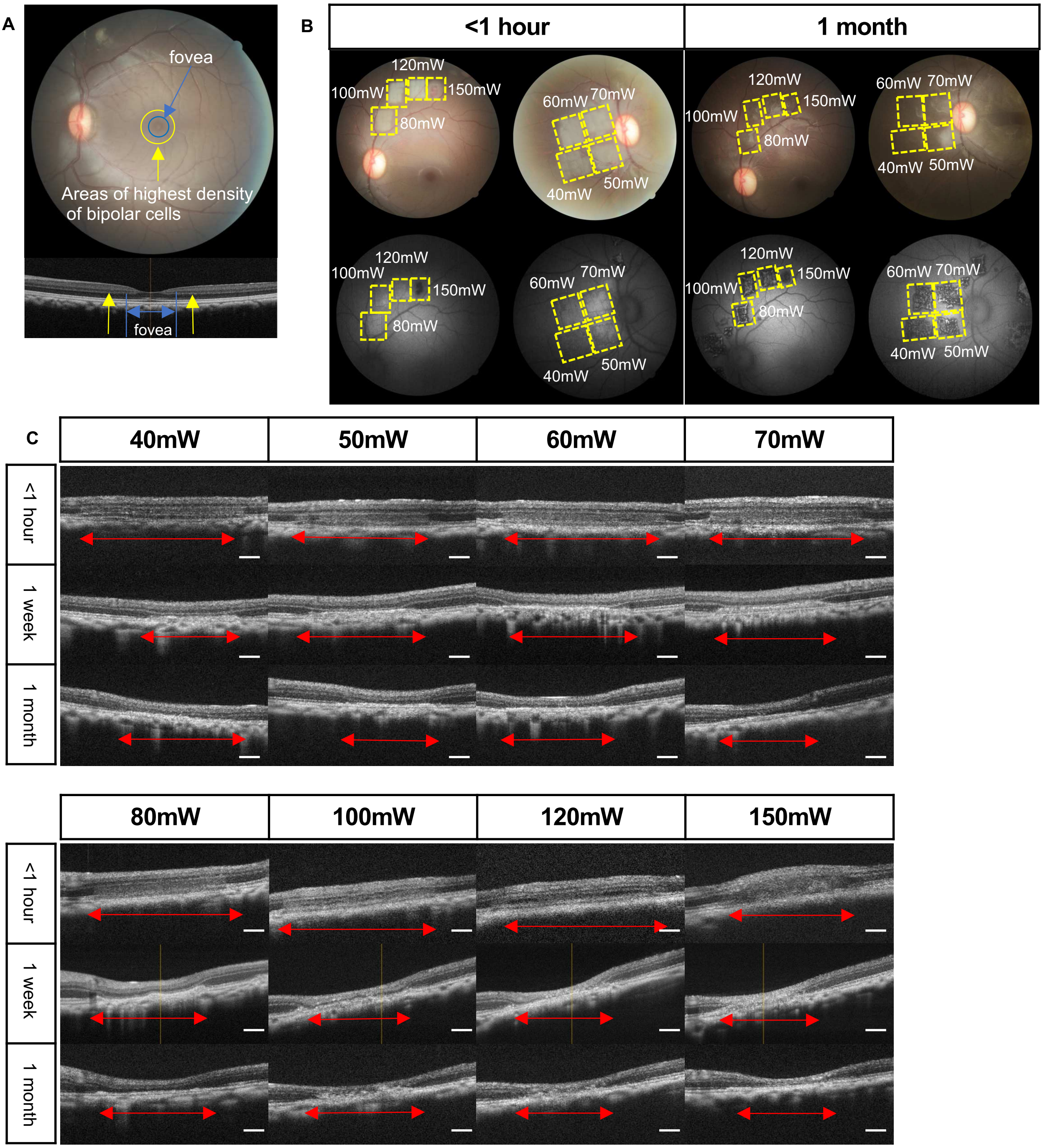


**Supplementary Figure 1. Optimization of laser power from 40mW to 150mW.** (A) The eccentricity range of fovea is from 0.13 to 0.6 mm (blue circle). The eccentricity of the highest density of BCs are 1mm (yellow circle). (B) Representative fundus images of 80-150mW (Upper left) and 40-70mW (Upper right) and autofluorescence images of 80-150mW (Lower left) and 40-70mW (Lower right) taken < 1 hour and 1 month after laser. (C)Sectional views of OCT imaging of 40-150mW < 1hour, 1 week, and 1 month after laser. The laser area was indicated by red two-headed arrow. Scale bars: 300μm (B)

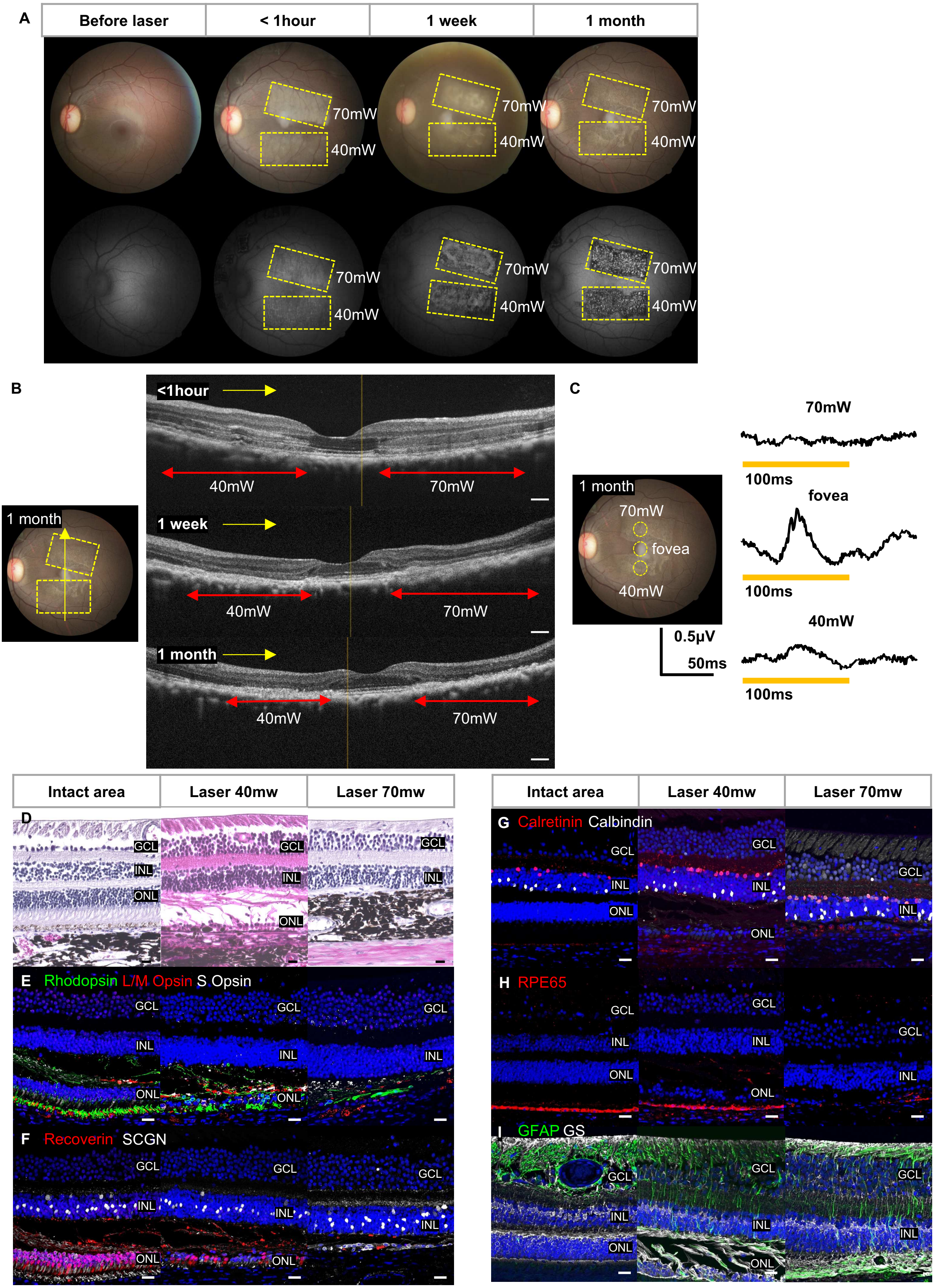
A
Before laser
< 1hour
1 week
1 month
70mW
40mW
B
1 month
<1hour
1 week
40mW
70mW
C
70mW
fovea
40mW
100ms
0.5μV
50ms
D
Intact area
Laser 40mw
Laser 70mw
GCL
INL
ONL
E Rhodopsin L/M Opsin S Opsin
F Recoverin SCGN
G Calretinin Calbindin
H RPE65
I GFAP GS

**Supplementary Figure 2. Laser-induced retinal degeneration monkey model involving para- and peri-fovea.** (A) Representative fundus images (Upper) and autofluorescence images (Lower) taken before laser treatment, <1 hour, 1 week, and 1 month after laser treatment. The upper area from the fovea was lasered at 70mW and the lower area at 40mW (yellow dot). (B) Cross-sectional OCT images of the yellow line on the fundus, taken <1 hour, 1 week, and 1 month after laser treatment. The left side of the OCT was at 40mW and the right side at 70mW (red two-headed arrow) (C) One month after laser, FMERG was stimulated at 70mW (upper), at the fovea (central), and at 40mW (lower) (yellow dot circle). (D-I) Histological images of retinas compared with intact area, laser 40mW, and laser 70mW. (D) Representative images of Hematoxylin &Eosin stained retinal sections showing ONL loss of Laser 70mW. (E-I) Retinal sections stained with Rhodopsin, L/M opsin and S opsin (E), Recoverin and SCGN (F), Calretinin and Calbindin (G), RPE65 (I), GFAP and GS (J). Scale bars: 300µm (B), 20µm (D), and 25µm (E-I)

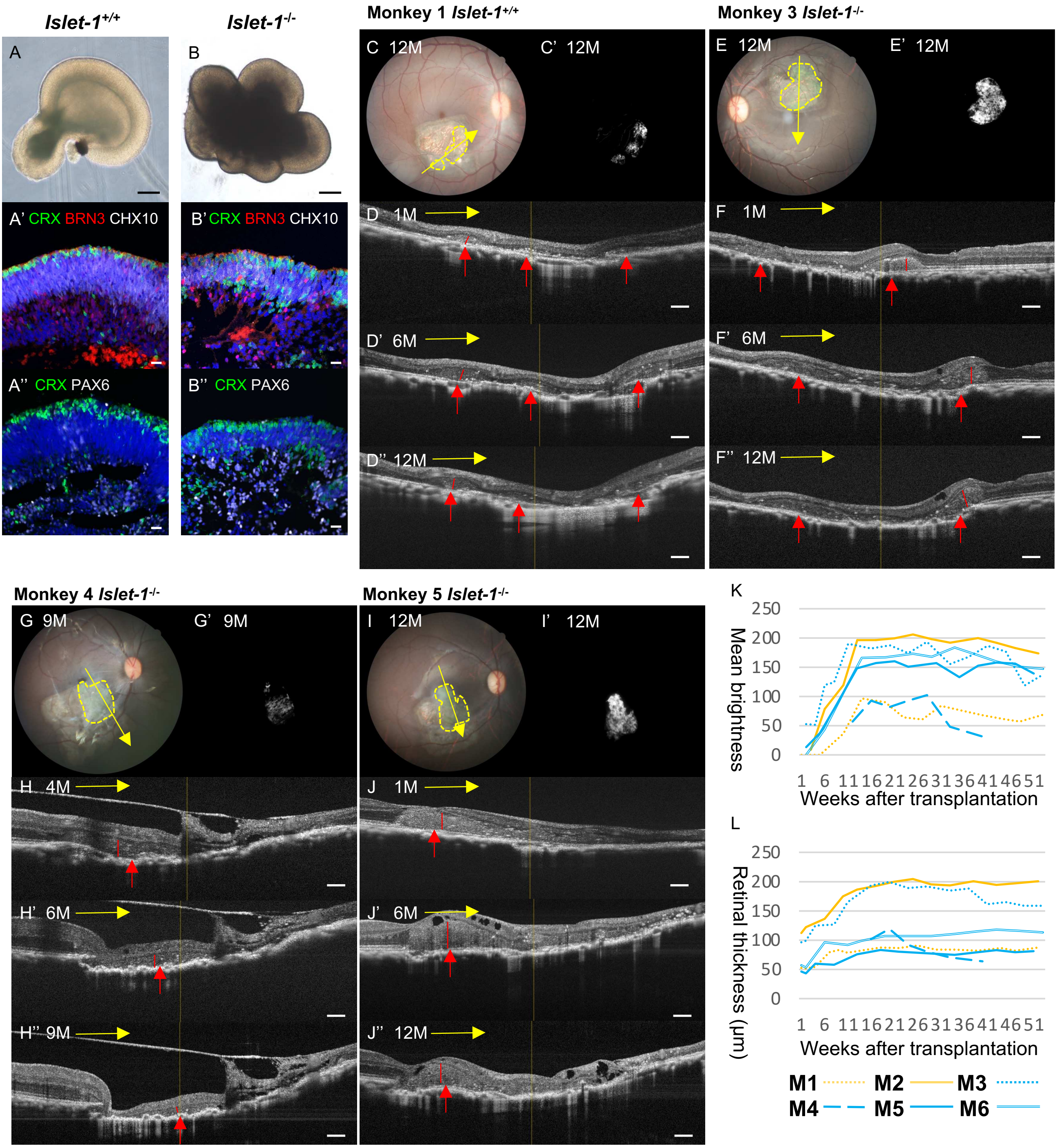


**Supplementary Figure 3. In vivo imaging of human ESC-derived RO sheets after transplantation.** (A, B) Bright field images of *Islet-1*$^{+/+}$ (A) and *Islet-1*$^{-/-}$ (B) human ESC ROs (approximately DD60). (A'), (B') Immunostaining of *Islet-1*$^{+/+}$ (A') and *Islet-1*$^{-/-}$ (B') human ESC retinal organoids for CRX, BRN3, and CHX10 (approximately DD60). (A'', B'') Immunostaining of *Islet-1*$^{+/+}$ (A'') and *Islet-1*$^{-/-}$ (B'') human ESC retinal organoids for CRX and PAX6 (approximately DD60) (C-J'') Representative fundus images (C, E, G, I), FA images of CRX::Venus (C', E', G', I'), and OCT images (D-D'', F-F'', H-H'', J-J'') of the transplanted grafts in the monkey eyes (Monkey 1, 3, 4, 5, respectively). (K) Line graph of temporal changes in Mean brightness of CRX::Venus for each transplanted eye. (L) Line graph of temporal changes in graft thickness for each transplanted eye. Scale bars: 100μm (A, B), 20μm (A'-A'', B'-B''), and 300μm (D-D'', F-F'',H-H'', J-J'')

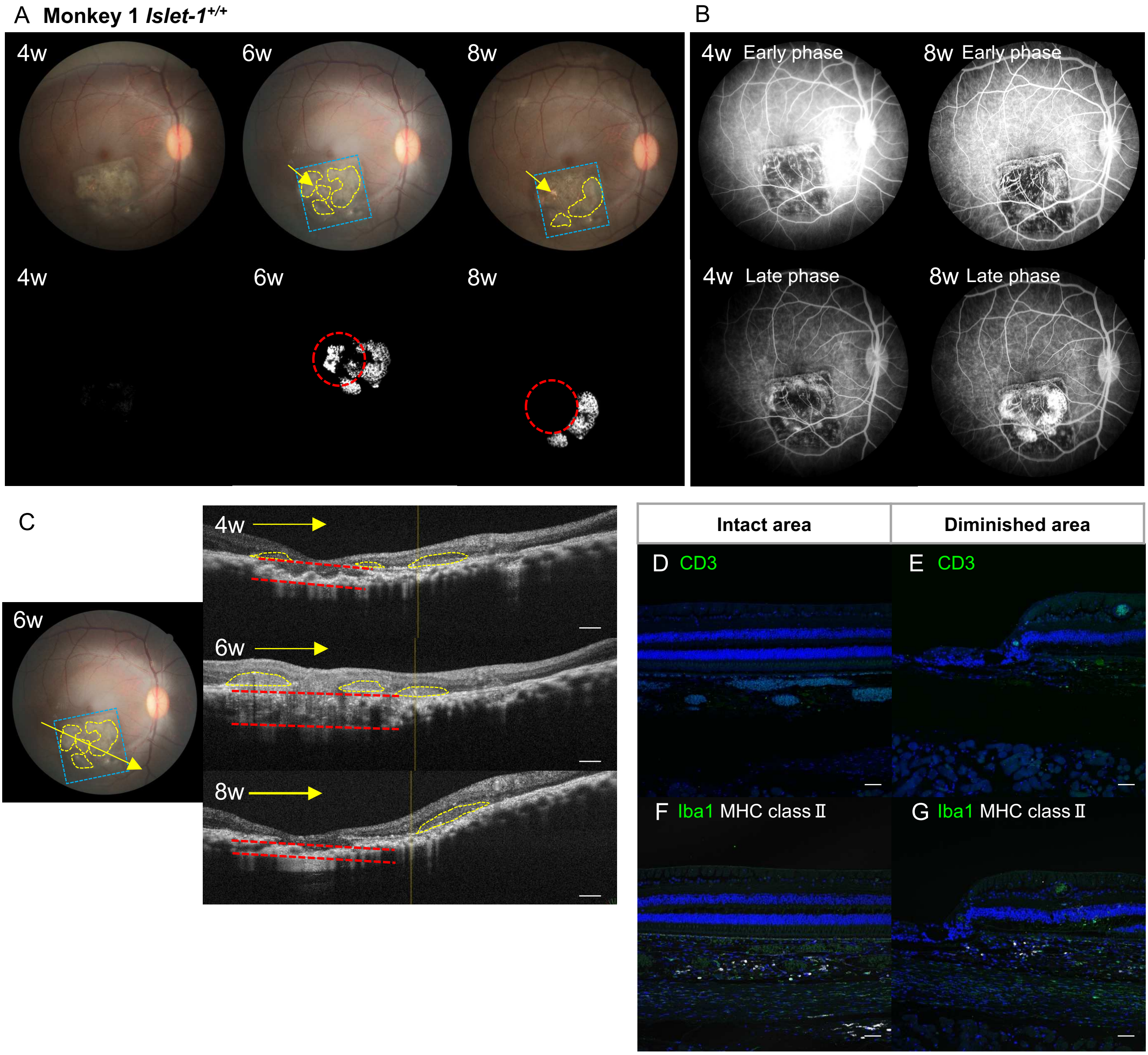


**Supplementary Figure 4. Evaluation of Monkey 1 before and after graft loss.** (A) Representative fundus images of the transplanted graft (yellow dot) before and after graft loss (red dot circle). Laser area was indicated by light blue dot, Insertion site was indicated by yellow arrow. (B) Representative FA images of the transplanted graft before (4w) and after (8w) graft loss. (C) Representative OCT images of the transplanted graft before (6w) and after (8w) graft loss. The grafts (yellow dot) were located at the laser area (light blue dot) Choroidal thickness was indicated by red dot line. (D-G) Retinal sections were stained with CD3 (D, E), Iba1 (F, G), and MHC class II (F, G) in the intact area and diminished area. Scale bars: 300µm (c), 50µm (d-g)

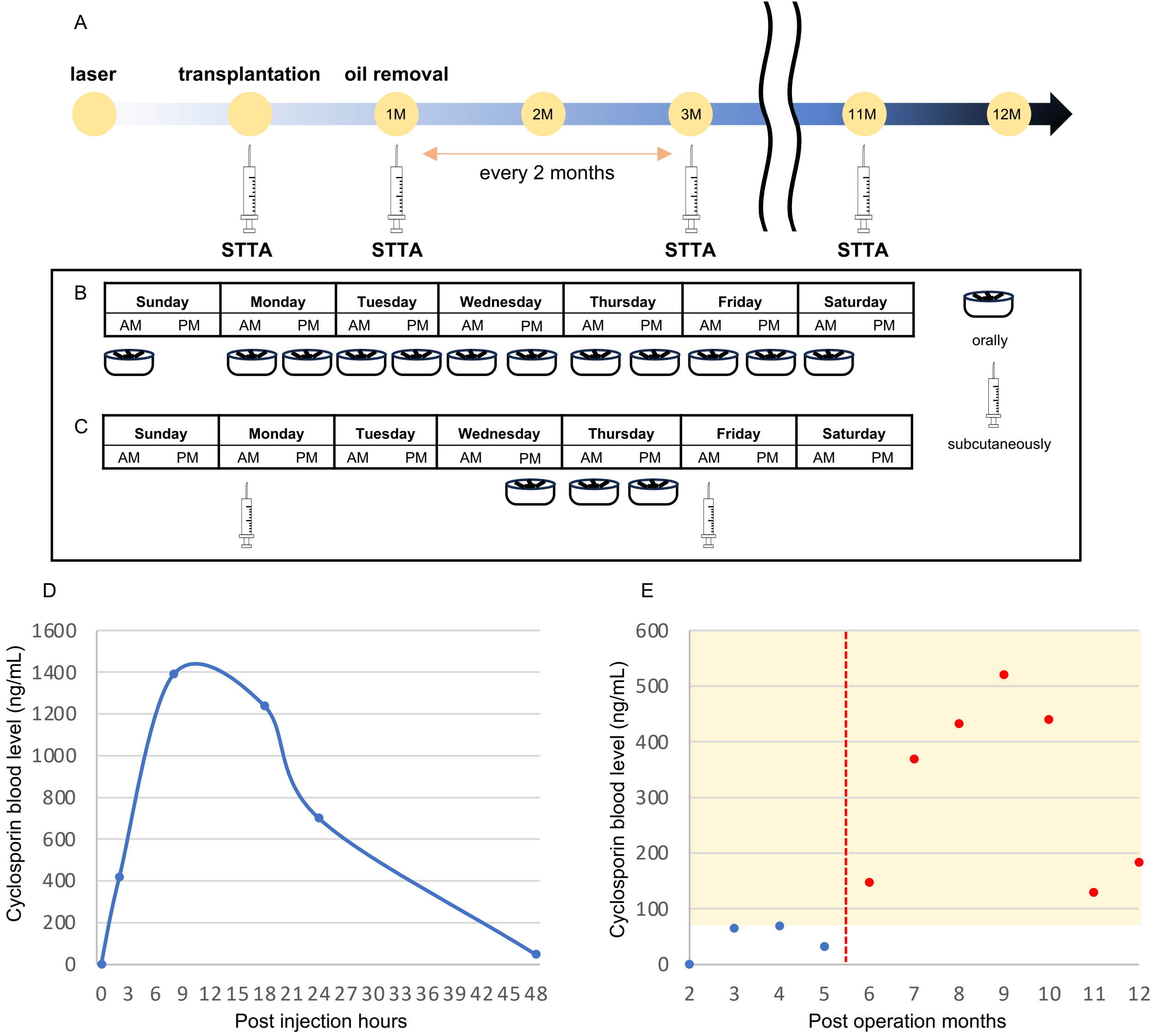


**Supplementary Figure 5. Management of local and systemic immunosuppression.** (A) Local immunosuppressant injection protocol. STTA was administered at the time of transplantation and oil removal, and then every 2 months. (B) Systemic immunosuppression protocol of cyclosporin orally. (C) Systemic immunosuppression protocol of oral and subcutaneous administration of cyclosporin (D) Cyclosporine blood concentration profile after subcutaneous administration of cyclosporin. Cyclosporine concentrations were measured over time in four monkeys. (E) Changes in blood concentration of cyclosporine in monkey 3. Blue dots represent cyclosporin managed by only orally, whereas red dots represent cyclosporin managed subcutaneously and orally.

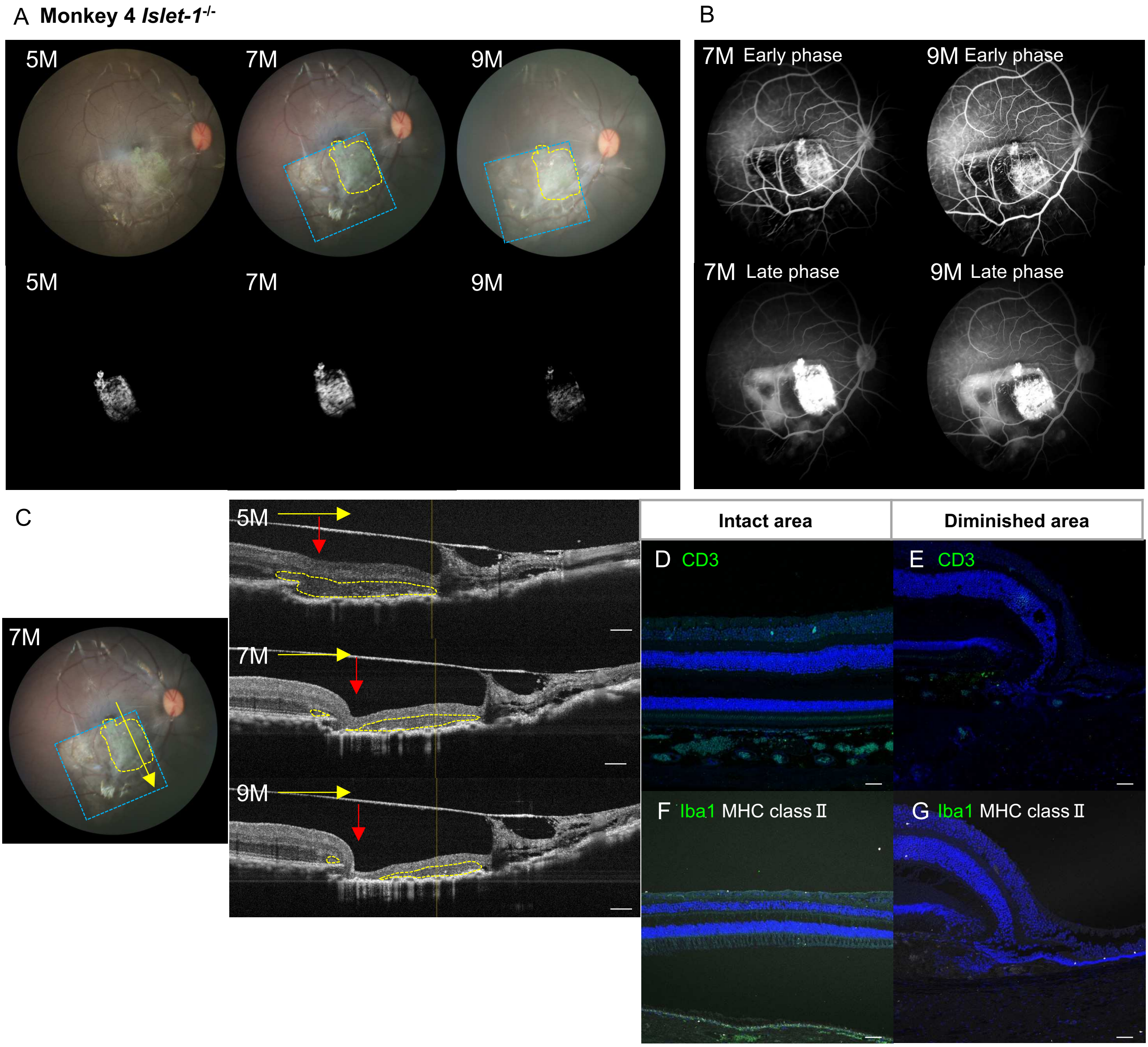


**Supplementary Figure 6. Evaluation of Monkey 4 before and after graft loss.** (A) Representative fundus images of the transplanted graft (yellow dot) before and after graft loss. (laser area indicated by light blue dot) (B) Representative FA images of the transplanted graft before (7M) and after (9M) graft loss. (C) Representative OCT images of the transplanted graft before (7M) and after (9M) graft loss. The grafts (yellow dot) were located at the laser area (light blue dot), and the diminished area was indicated by red arrow. (D-G) Retinal sections were stained with CD3 (D, E), Iba1 (F, G), and MHC class II (F, G) in the intact area and diminished area. Scale bars: 300µm (c), 50µm (d-g)

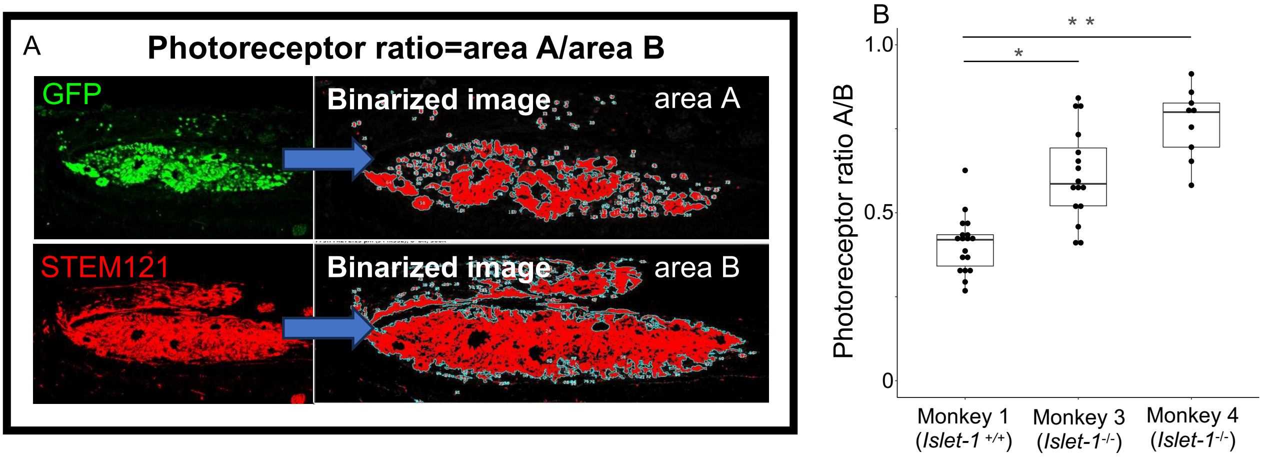


**Supplementary Figure 7. The ratio of photoreceptor between the *Islet-1*$^{+/+}$ and *Islet-1*$^{-/-}$ grafts**
(A) Schematic representations of PRCs ratio with binarized images of GFP (graft PRCs) and STEM121 (graft cells). The PRC ratio was defined as the area of GFP divided by the area of STEM121. (B) The PRC ratio in the grafts of each monkey (monkey 1: 18 sections, monkey 3: 16 sections, monkey 4: 9 sections). The PRC ratio of monkey 3 and monkey 4, which were transplanted with *Islet-1*$^{-/-}$ retinal organoid sheets, was significantly higher than that of monkey 1 (*p = 2.427e-05, **p=8.535e-07, Wilcoxon rank sum test).

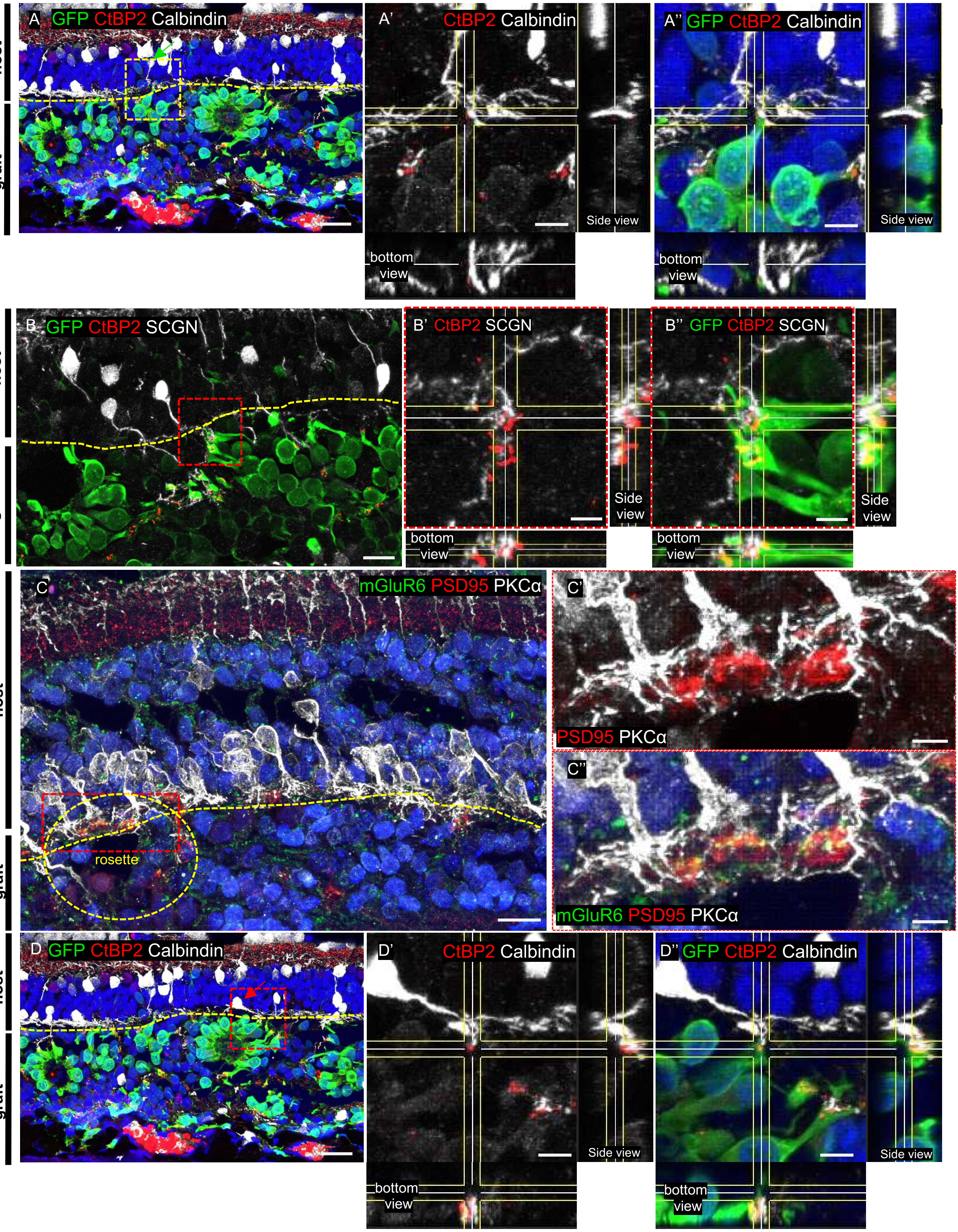
A GFP CtBP2 Calbindin
host
graft
A' CtBP2 Calbindin
Side view
bottom view
A'' GFP CtBP2 Calbindin
Side view
bottom view
B GFP CtBP2 SCGN
host
graft
B' CtBP2 SCGN
Side view
bottom view
B'' GFP CtBP2 SCGN
Side view
bottom view
C mGluR6 PSD95 PKCα
host
graft
rosette
C'
PSD95 PKCα
C''
mGluR6 PSD95 PKCα
D GFP CtBP2 Calbindin
host
graft
D' CtBP2 Calbindin
Side view
bottom view
D'' GFP CtBP2 Calbindin
Side view
bottom view

**Supplementary Figure 8. Host-graft synaptic formation of transplanted PRCs.** (A-B'') Presynaptic marker CtBP2 was localized on the margin of the GFP (graft PRCs) and the dendrite tips of calbindin (A-A'') and SCGN (B-B'')-positive BCs. (C-C'') Presynaptic marker PSD95 and postsynaptic marker mGluR6 were present at the dendritic tips of PKCα-positive BCs. (D-D'') Presynaptic marker CtBP2 was localized on the margin of the GFP (graft PRCs) and the dendrite tips of Calbindin-positive horizontal cells. Scale bars: 15μm (A, B, C, D) 4μm (A', A'', B', B'', D', D'') 5μm (C', C'')

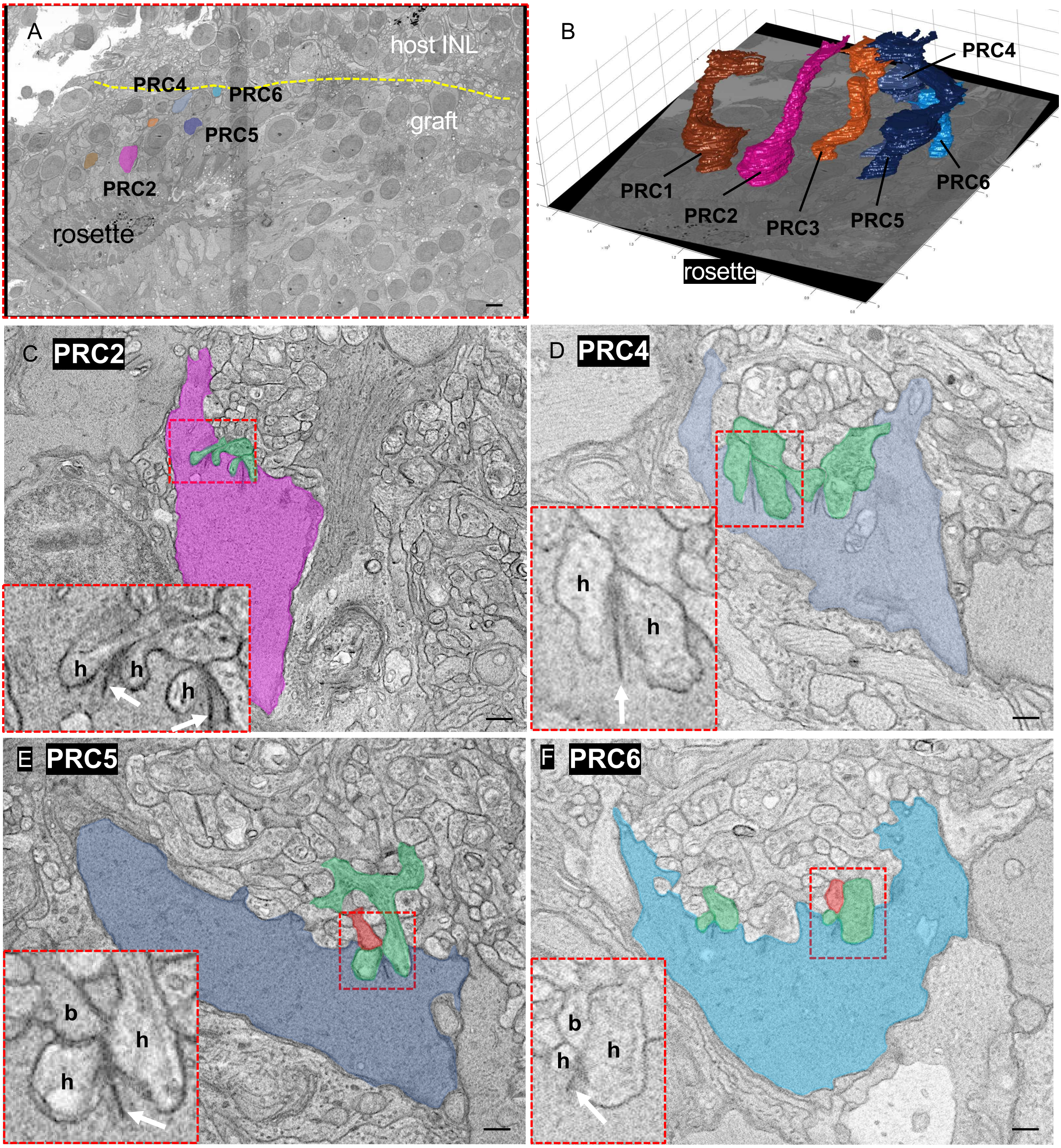


**Supplementary Figure 9. Synaptic connections between host BCs and graft PRCs observed in EM.** (A) Enlarged view of the area within the red dotted rectangle in (Figure 7C), showing the rosette that includes each traced PRC. PRC2: pink (C), PRC4: blue (D), PRC5: dark blue (E), PRC6: light blue (F). (B) 3D reconstructed image of graft photoreceptors extending from the rosette into the host INL; six PRCs were traced. (C-F) EM images of the axon terminals of individual PRCs. white arrows showed synapse ribbon. b, bipolar cell; h, horizontal cell scale bars: 5μm (A) 500nm (C-F)

**Supplementary Table 1. Summary of transplanted eyes of monkeys**

| Monkey number | M1 | M2 | M3 | M4 | M5 | M6 |
|---|---|---|---|---|---|---|
| Age at transplantation | 6 | 3 | 3 | 3 | 3 | 9 |
| Operated eye | Right | Right | Left | Right | Right | Right |
| Organoid type | Wild type | Wild type | *Islet1*$^{-/-}$ | *Islet1*$^{-/-}$ | *Islet1*$^{-/-}$ | *Islet1*$^{-/-}$ |
| Differentiation day of transplantation | 56 | 63 | 63 | 59 | 63 | 64 |
| Transplantation day after laser | 29 | 43 | 18 | 43 | 47 | 28 |
| Postsurgical events | - | - | - | Postoperative endophthalmitis | Cataract | - |
| Graft disappearing | + | - | - | + | - | - |
| Histological evaluation day after transplantation | 427 | - | 413 | 286 | 402 | - |

| **Supplementary Table 2. Summary of primary antibody** | | |
|---|---|---|
| Goat polyclonal anti-BRN3 | Santa Cruz biotechnology | Cat#sc-6026 RRID: AB_673441 |
| Rabbit polyclonal anti-Calbindin | Millipore | Cat#ABN2192 RRID: AB_2935805 |
| Mouse monoclonal anti-Calretinin | Millipore | Cat#MAB1568 RRID: AB_94259 |
| Rabbit polyclonal anti-CD3 | Abcam | Cat#ab5690 RRID: AB_305055 |
| Mouse monoclonal anti-CHX10 | Santa Cruz biotechnology | Cat#sc-365519 RRID: AB_10842442 |
| Mouse monoclonal anti-CtBP2 | BD | Cat#612044 RRID: AB_399431 |
| Rabbit polyclonal anti-EAAT2 | Proteintech | Cat#22515-1-AP RRID: AB_2879112 |
| Rabbit polyclonal anti-GFAP | DAKO | Cat#Z0334 RRID: AB_ 10013382 |
| Rabbit polyclonal anti-GFP | abcam | Cat#ab290 RRID: AB_ 2313768 |
| Mouse monoclonal anti-GS | Millipore | Cat#MAB302 RRID: AB_ 2110656 |
| Rabbit polyclonal anti-Iba1 | Wako | Cat#019-19741 RRID: AB_839504 |
| Rabbit polyclonal anti-mGluR6 | Novus Biologicals | Cat#NLS4655 RRID: AB_343723 |
| Mouse Monoclonal anti-MHC classII | DAKO | Cat#M0775 RRID: AB_2313661 |
| Goat polyclonal anti-OPN1SW | Santa Cruz biotechnology | Cat#sc-14363 RRID: AB_2158332 |
| Rabbit polyclonal anti-Opsin Re＋G | Sigma-Aldrich | Cat#AB5405 RRID: AB_177456 |
| Rabbit polyclonal anti-Opsin Blue | Millipore | Cat#AB5407 RRID: AB_304867 |
| Mouse monoclonal anti-PAX6 | BD Biosciences | Cat#BD561462 RRID: AB_10715442 |
| Rabbit polyclonal anti-Peripherin-2 | Proteintech | Cat#18109-1-AP RRID: AB_10665364 |
| Anti-PDE6H Mouse monoclonal | Sigma-Aldrich | Cat# sc-398478 |
| Mouse monoclonal anti-PKCα | Sigma-Aldrich | Cat#P5704 RRID: AB_477375 |
| Mouse monoclonal anti-PSD95 | Biolegend | Cat#810401 RRID: AB_2564750 |
| Rabbit polyclonal anti-Recoverin | Proteintech | Cat#10073-1-AP RRID: AB_2178005 _ |
| Mouse monoclonal anti-Rhodopsin | Sigma-Aldrich | Cat#O4886 RRID: AB_260838 |
| Rabbit polyclonal anti-RPE65 | Takara | Cat#AS2352 |
| Sheep polyclonal anti-secretagogin | BioVendor R&D | Cat#RD184120100 RRID: AB_2034062 |
| Mouse monoclonal anti-STEM121 | Takara | Cat#Y40410 RRID: AB_ 2801314 |
